# Engineering Carbon Nanotube Quantum Well Defects with Recognition Tripeptides for Optical Detection of Extracellular Vesicles in Plasma

**DOI:** 10.64898/2026.06.01.729398

**Authors:** In-Jun Hwang, Junhee Kim, Aarti Patel, Leilei Zhang, Jacob Miller, Stanislav Piletsky, Cassandra L. Clift, Colin Hisey, YongJoo Kim, Mijin Kim

## Abstract

Extracellular vesicles (EVs) carry molecular signatures of their originating cells and have thus emerged as promising biomarkers. However, their clinical utility remains limited due to their low abundance and the modest sensitivity of current EV detection methods in complex biological environments. Here, we present a quantum well defect functionalized carbon nanotube sensor coupled with integrin-recognition RGD tripeptide for EV detection in human plasma. Leveraging the abundance of integrins on EV surfaces, we targeted α5β1, αVβ1, and αVβ3 subtypes. The nanosensor exhibited robust hypsochromic shifts in defect emission upon integrin binding, achieving sub-picomolar detection limits for integrin subunits and quantifying EVs at concentrations as low as 10^4^ EVs·mL^−1^ for glioblastoma, ovarian cancer, and fibroblast cell-derived EV types. Molecular dynamics simulation indicated that integrin docking at the RGD-coupled quantum defect can substantially reshape the interfacial environments of the quantum defects, explaining the high sensitivity in EV detection in complex biological media. Finally, transmembrane protein analysis validated the expression of surface integrins across the tested EV types. The modular nanosensor construct can be targeted to detect disease-associated EV subpopulations, advancing EV-based diagnostics.

## Introduction

Liquid biopsies have significant potential in precision oncology by providing dynamic molecular information about tumors through minimally invasive procedures.^1^ Extracellular vesicles (EVs) have emerged as promising targets within the field of liquid biopsy. EVs are nanoscale particles, secreted by cells and enriched with cellular constituents that mirror the molecular states of the originating cells ^2^. Tumor cell-derived EV profiling thus can yield comprehensive and actionable clinical insights. EV proteomic profiling, for instance, has achieved high diagnostic specificity by targeting tumor-specific antigens and identified tumor origins or subtypes based on protein signatures.^3^ EV-derived mRNA analysis can provide complementary information, including somatic mutations and resistance-associated transcripts.^4^ Collectively, these advantages position EVs as a promising source of cancer biomarkers, augmenting other liquid biopsy approaches such as circulating tumor DNA and protein panels.

Despite the unique advantages of EVs as cancer biomarkers, their clinical potential remains underutilized, in part due to technical challenges in detection. Tumor-derived EVs constitute a small fraction (<5%) of total circulating EVs^5^. Conventional isolation methods, such as ultracentrifugation and precipitation, are labor-intensive, require large sample volumes, and often compromise EV integrity, hindering reliable quantification.^6^ Thus, innovative molecular techniques that detect EVs from low sample volumes with high sensitivity and specificity are essential to harness the full potential of EVs.

Quantum well defects (QWDs) are synthetic functional groups introduced onto semiconducting single-walled carbon nanotubes via controlled covalent chemistry.^7^ Quantum defects create new fluorescence bands 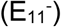 ∼100-300 meV redshifted from the intrinsic nanotube emission (E_11_).^8^ The defect fluorescence imparts new chemical sensitivities to quantum defect-modified nanotubes (QWNTs), making quantum defects the molecular focal point for chemical detection.^9-12^ Molecular recognition elements that have been successfully implemented in other sensing platforms^13-15^ may be adapted to QWNT systems, offering a systematic approach to optimize defect functionalization and sensitization for biochemical sensing applications.

Herein, we report the development of a QWNT-based nanosensor coupled with integrin-recognition RGD tripeptide to detect EVs in complex plasma environments. Given the abundance and RGD-binding properties of surface integrins on EVs, we selected α5β1, αVβ1, and αVβ3 integrin subtypes as our targets. Quantum defect sensitization was achieved through bioconjugation of RGD peptides to carboxyl aryl QWDs. The nanosensor demonstrated robust hypsochromic shifts in the quantum defect emission upon exposure to integrin subunits in plasma. The sub-picomolar level detection limits outperformed conventional immunoassays by four orders of magnitude. Molecular dynamics simulations indicate that the RGD-coupled defect site stably engages the integrin-binding pocket, providing a structural rationale for integrin-specific emission response. The nanosensor was able to detect ovarian cancer (OVCAR4), glioblastoma (A172), and fibroblast (HDFa)-derived EVs as low as 10^4^ EVs·mL^-1^. Notably, the nanosensor detected surface integrins on fibroblast-derived EVs, a population for which EV surface integrin biology remains poorly characterized. EV surfaceome proteomics analysis confirmed the presence of surface integrins, providing a molecular basis for the observed sensor responses. Our study suggests that RGD-QWNT nanosensors can serve as tools for probing integrin-carrying EV subpopulations, including those from stromal cell types. The modular design of RGD-QWNTs can be applied to diverse short peptides to detect disease-associated EVs and monitor changes in EV surface composition in biological media.

## Results

### Synthesis of RGD-QWNTs

RGD tripeptide (Arg-Gly-Asp) is a short amino acid sequence that serves as a recognition motif for integrins in the αV, β1, and β3 families. Within the tripeptide, the Arg residue forms salt bridges with the α-subunit, the Asp residue coordinates a divalent cation at the β-subunit metalion-dependent adhesion site, and the Gly residue contributes via weak hydrogen bonding, ensuring the specificity and stability of the protein-tripeptide interaction^16-19^. To sensitize carbon nanotubes to integrins and EVs, we developed an RGD tripeptide-QWNT construct (**Fig. 1A**).

**Figure 1.**
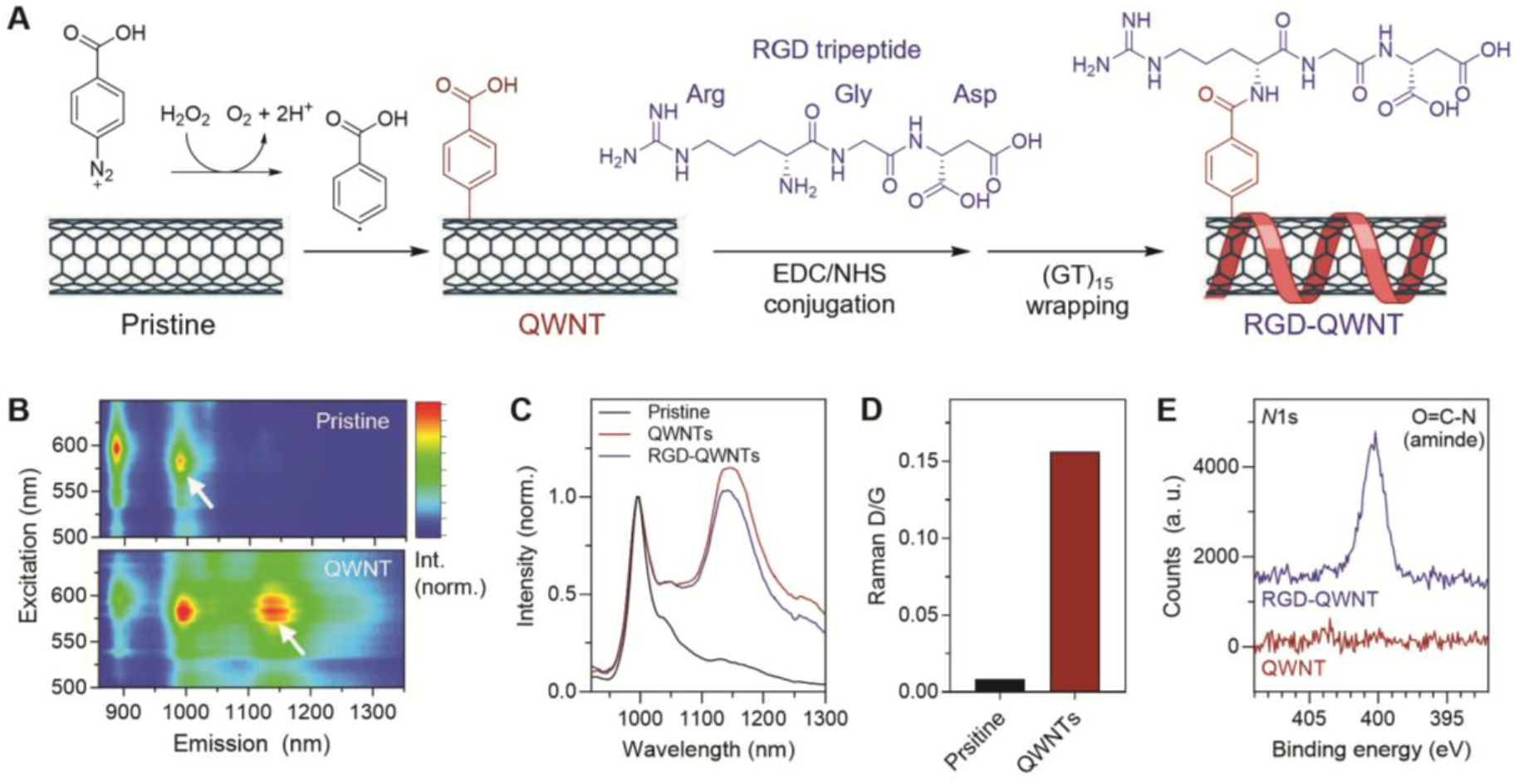
Design and characterization of RGD-QWNTs. (**A**) Schematic illustration of the RGD-QWNT synthesis from a pristine carbon nanotube. (**B**) Excitation-emission fluorescence maps of pristine nanotubes and QWNT. Arrows indicate the intrinsic E_11_ and defect-induced 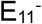 emission bands. (**C**) Fluorescence emission spectra of pristine, QWNTs, and RGD-QWNTs. (**D**) Raman D-band to G-band ratio of pristine and QWNTs. (**E**) *N* 1s XPS spectra showing the appearance of the amide (O=C-N) peak after RGD conjugation.

The (6,5) chirality carbon nanotubes were covalently functionalized with 4-carboxyaryl quantum well defects via solid-state diazonium chemistry (see Methods). To confirm the carboxyaryl groups covalently attached to the nanotubes, we characterized the nanotubes with fluorescence and Raman spectroscopy. For short-wave near infrared (SWIR) fluorescence characterization, the QWNTs were dispersed in an aqueous solution of sodium deoxycholate (see Methods). The SWIR excitation-emission fluorescence map of the QWNTs showed the intrinsic emission band of (6,5) carbon nanotubes (E_11_) at 996 nm and a bright new emission band 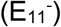 at 1,143 nm, resonant to (6,5) excitation profile (**Fig. 1B-C**). We optimized the intensity ratio between E_11_ and 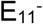 of QWNTs to 1:1 to 1:2 to ensure that both emission bands were visible, enabling the observation of subtle changes in emission wavelength. The Raman spectroscopy of the QWNTs exhibited an increased disorder-induced D band at 1315 cm^-1^ with respect to the in-plane stretching mode of the sp^2^ carbon lattice, G-band at 1,590 cm^-1^, indicating the increase of sp^3^ carbons (**Fig. 1D**, S1).

RGD was then bioconjugated to solid-state QWNTs via a carbodiimide crosslinker chemistry. The primary amine of arginine in the RGD sequence specifically reacts with the carboxyl group on the quantum well defect to form a stable amide bond. To confirm the covalent coupling of RGD to the carboxylaryl quantum well defect, we characterized the QWNTs using X-ray photoelectron spectroscopy (XPS). RGD-QWNTs exhibited an *N* 1s characteristic peak at 400.3 eV, corresponding to the amide bonds of RGD, whereas QWNTs showed no noticeable peaks in the *N* 1s region (**Fig. 1E**). The amide C-N and O=C-N bond of the RGD tripeptide newly appeared at 286.2 and 288.3 eV, respectively, in the C 1s XPS spectrum of RGD-QWNTs (Fig. S2).

To confer colloidal stability of the QWNTs in aqueous solution, the RGD-QWNTs were wrapped with a single-stranded DNA oligonucleotide (GT)_15_ (see Methods). The absorption profile of QWNTs remained the same after RGD conjugation (Fig. S3), suggesting that the RGD conjugation did not substantially disturb the sp^2^ carbon lattice of the nanotube and preferentially conjugated to the quantum well defects. While RGD coupling to QWNTs did not change the E_11_ emission profile, the defect-induced 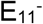 bands blue-shifted by 4 nm, and the E_11_ to 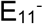 intensity ratio was slightly reduced from 1.15 to 1.03 (**Fig. 1C**, S4). This change is presumably due to the fact that converting the terminating moiety of aryl defects from a carboxylic to an amide group makes the quantum well defects more electron-donating, as predicted by the Hammett constant, resulting in an increase in the defect state energy levels^20^.

### Fluorescence response of RGD-QWNTs to integrins

We investigated the fluorescence responses of RGD-QWNTs to three representative integrin subunits, α5β1, αVβ1, and αVβ3 in 20% human plasma. RGD-QWNTs were titrated with integrins at concentrations ranging from 0.001 to 100 ng·mL^-1^ (**Fig. 2A-B**, S5). The spectral response of the RGD-QWNTs saturated within 60 min of incubation (**Fig. 2C**). All three integrin subunits induced a concentration-dependent spectral shift of the 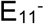 band. At the highest integrin concentration, 100 ng·mL^-1^, the 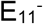 band blue shifted by 1.5 nm, whereas the E_11_ wavelengths remained unchanged (**Fig. 2D**, S6). The limits of detection (LOD) of RGD-QWNTs for integrins were determined to be 2.6, 6.3, and 2.4 pg·mL^-1^ for α5β1, αVβ1, and αVβ3, respectively.

**Figure 2.**
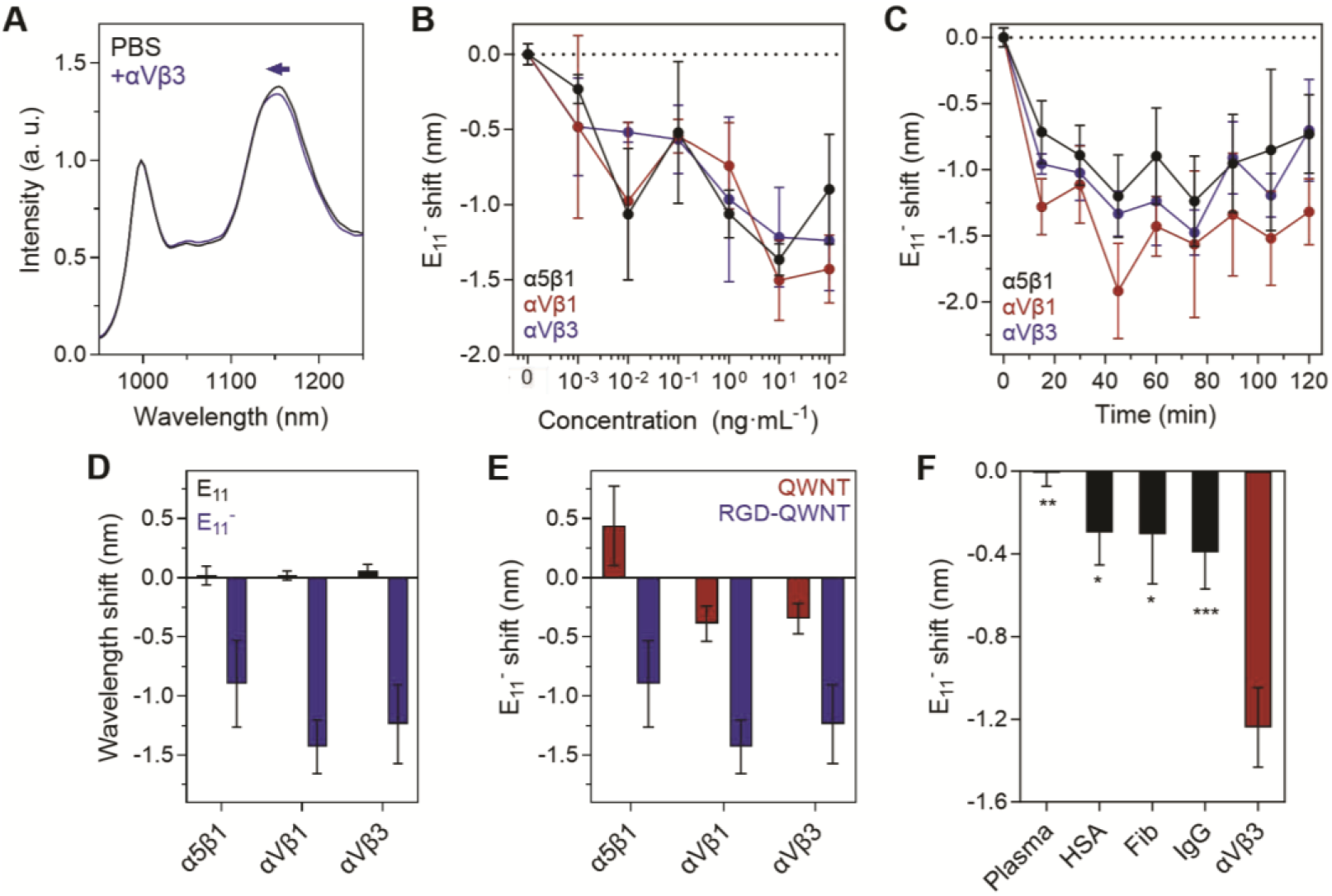
Response of RGD-QWNTs to integrin in 20% human plasma. **(A)** Fluorescence spectra after 2-hour incubation in addition of PBS (black) or integrin αVβ3 (100 ng**·**mL^-1^) (blue). (**B**) Concentration-dependent 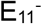 peak shift at 2 hour timepoint. (**C**) 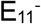 peak shift as a function of incubation time after the addition of integrin subunits (100 ng**·**mL^-1^). (**D**) Wavelength shift of E_11_ (black) and 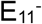 (blue) peaks after incubation of integrin subunits at 60 min (100 ng**·**mL^-1^). (**E**) 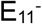 shift of QWNTs and RGD-QWNTs after 60 min of incubation with integrin subunits. (**F**) 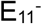 shift in the presence of blood-abundant proteins. All error bars represent standard deviation from the mean values (n = 3). *p < 0.05; **p < 0.01; ***p < 0.001 (n = 3) in t-test.

To assess the effects of RGD tripeptide conjugation on the sensitivity, we compared the spectral responses of QWNTs and RGD-QWNTs. Upon addition of 100 ng·mL^-1^ integrins, QWNTs did not induce statistically significant spectral shifts (**Fig. 2E**). The specificity of RGD-QWNTs was further evaluated against abundant plasma proteins, including human serum albumin (HSA), fibronectin (Fib), and immunoglobulin G (IgG) at 100 ng·mL^-1^ (**Fig. 2F**). These proteins induced significantly smaller 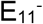 wavelength shifts compared to integrin αVβ3, confirming the selectivity of RGD-QWNTs.

### Molecular dynamics simulation of integrin binding to RGD-QWNTs

To investigate the molecular basis of integrin binding to RGD-QWNTs, we performed molecular dynamics (MD) simulations using integrin αVβ3 as a model receptor (**Fig 3**, see Methods). To isolate the effects of the RGD moiety, we compared RGD-conjugated QWNT with control QWNT with a carboxyl aryl defect. In the post-equilibrium window (400-800 ns), RGD-QWNTs formed significantly more defect-integrin contacts than the control (51.6 ±5.5 vs. 4.6±4.1 atoms within 3.5 Å interfacial distances, Fig S7), indicating stable, QWD-mediated engagement. Contact frequency maps projected onto the integrin surface revealed distinct binding geometries and retention between the two constructs. The RGD-conjugated QWD maintained persistent contacts within the canonical RGD-binding pocket of integrin αVβ3 (**Fig 3A**). The high contact frequency region coincided with the known RGD-recognition site. In contrast, the control QWNT transiently engaged through the nanotube sidewall.

**Figure 3.**
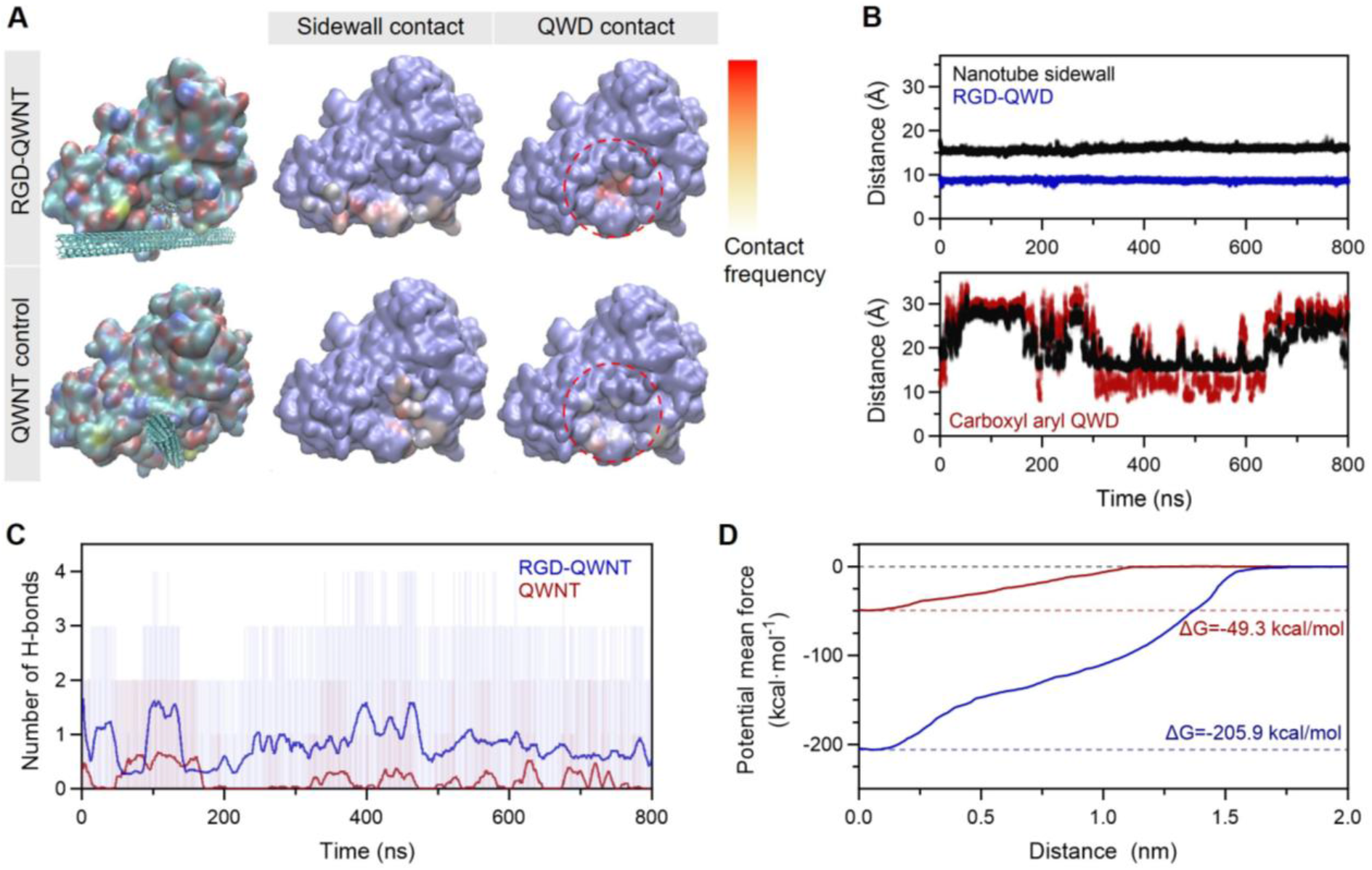
Molecular dynamics simulations of integrin αVβ3 interaction with QWNTs. (**A**) Contact frequency maps superimposed onto the integrin αVβ3 surface for RGD-QWNT (top) and control QWNT (bottom) within the post-equilibrium window (400-800 ns). Left: Integrin-QWNT complex; middle: sidewall-integrin contacts; right: QWD-integrin contacts. Red dashed circle indicates the canonical RGD-binding pocket. (**B**) Center-of-mass distance between Metal Ion-Dependent Adhesion Site (Mg^2+^) and the nanotube sidewall (black) or QWDs (blue/red) for RGD-QWNT (top) and control (bottom). (**C**) Number of hydrogen bonds between the integrin and QWNTs. (**D**) Potential mean force profiles for integrin-QWNT unbinding as a function of separation distance. Corresponding binding free energies (ΔG) are indicated.

We then investigated the stability of integrin-QWNT interactions. The center-of-mass (COM) distance between the Metal Ion-Dependent Adhesion Site (MIDAS, Mg^2+^) of the integrin^21^ and the quantum well defect was tightly distributed at 8.7±0.2 Å for RGD-QWNT throughout the post-equilibrium window while control QWNT drifted between 7.6-35.4 Å with standard deviation of 7.6 Å (**Fig. 3B**). Consistently, RGD-QWNTs maintained 1-4 hydrogen bonds between the RGD-conjugated quantum well defect and β3 pocket residues while the hydrogen bonding in control QWNT collapsed at the QWD escaped the pocket (**Fig. 3C**). At the residue level, root-mean-square fluctuation (RMSF) suppression in RGD-QWNT was localized to β3 residues that directly contact the RGD motif, Asn215, Tyr122, Ser121, Ser123, and Glu220, consistent with stable recognition at the RGD-coupled defect (Fig. S8). Residue-resolved COM distances confirmed the canonical RGD-β3 contacts throughout the trajectory (Fig. S9). In control QWNT, RMSF suppression appeared at αV subunit residues, Asp150 and Tyr178 (Fig. S10), reflecting nonspecific steric contact by the nanotube sidewall rather than specific recognition.

The overall binding affinity was assessed by computing the potential of mean force along the QWD-integrin separation coordinate (**Fig. 3D**). RGD-QWNT exhibited a binding energy of - 205.89 kcal·mol^-1^, compared to -49.33 kcal·mol^-1^ for control QWNT (ΔG = 156.56 kcal·mol^-1^ in favor of RGD-QWNT). Interaction energy decomposition indicated that favorable RGD-defect-integrin interactions are dominated by electrostatic contributions localized at the defect-integrin interface, with smaller but non-negligible van der Waals contributions (Fig. S11). On the other hand, the defect-integrin interaction in the control QWNT was carried by nonspecific sidewall contacts (Table S1). This is consistent with canonical RGD-integrin recognition, in which the Asp carboxylate of the RGD motif coordinates the MIDAS metal ion, stabilized by hydrogen bonds with β3 residues Tyr122 and Asn215.^19^ More importantly, the electrostatically driven binding interactions explain that the integrin selectively perturbs the local dielectric environment of the QWD and modulates the 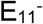 emission observed experimentally in Fig 2.

### Fluorescence response of RGD-QWNTs to extracellular vesicles

We validated the functionality of RGD-QWNTs using EVs derived from human fibroblasts (HDFa), glioblastoma (A172), and ovarian cancer (OVCAR4) bioreactors. These EVs represent physiologically distinct origins, allowing evaluation of RGD-QWNT performance with diverse clinical implications. HDFa, A172, and OVCAR4 cell lines express integrins such as α5β1, αVβ1, and αVβ3, and A172 and OVCAR4-derived EVs are known to carry specific integrin heterodimers on their surfaces. Thus, the presence of integrins on EV surfaces can confer specific EV-nanotube interactions (**Fig. 4A**). To test this, we first isolated EVs from bioreactors via ultracentrifugation, size exclusion chromatography, and filtration (Fig. S12). The size and morphology of EVs were characterized using nanoparticle tracking analysis (NTA) and negative staining TEM. The mean particle sizes of HDFa, A172, and OVCAR4-derived EVs were 80.0 ± 52.8, 48.9 ± 47.1, and 96.1 ± 67.6 nm, respectively (**Figs. 4B**, S13). The TEM images showed morphologically heterogeneous, sub-100 nm sized EVs (Fig. S14). EV identity is supported by enrichment of the tetraspanin CD81 and cytosolic GAPDH and depletion of the ER marker, GRP94 in isolated EVs relative to parental cell lysates^22^ (Fig. S15).

**Figure 4.**
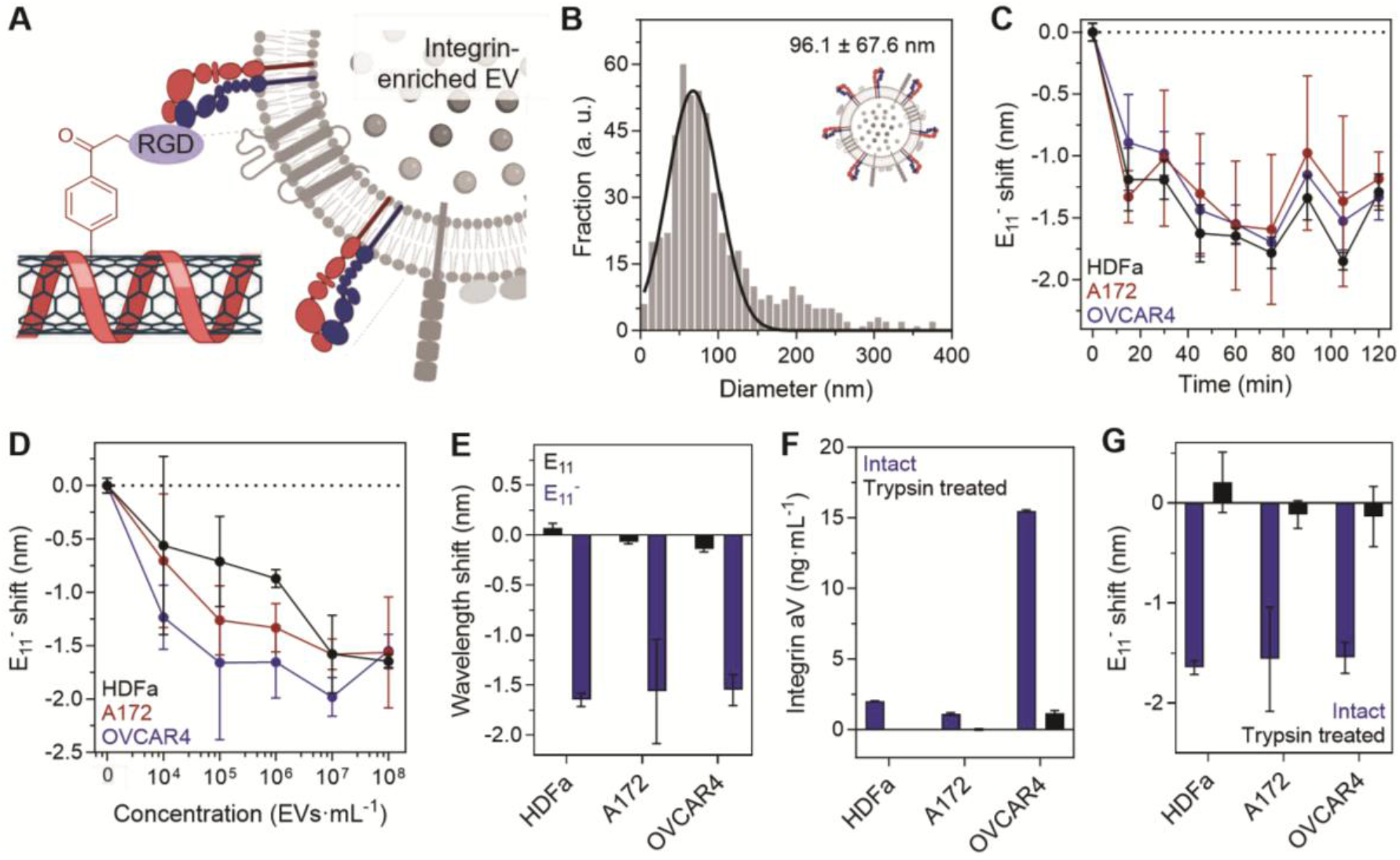
Response of RGD-QWNTs to extracellular vesicles in 20% human plasma. **(A)** Schematic illustration of quantum well sensitization of RGD-QWNTs. (**B**) Hydrodynamic size of OVCAR4. (**C)** Time-dependent 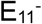 peak shift after addition of EVs (1×10^8^ EVs·mL^-1^). (**D**) 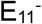 peak shift as a function of concentration EVs. **(E)** Wavelength shift of E_11_ (black) and 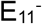 (blue) peaks after incubation of EVs for 60 min (1×10^8^ EVs·mL^-1^). **(F)** Quantification of integrin αV after shaving of proteins of EVs (1×10^12^ EVs·mL^-1^) by ELISA. **(G)** 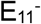 shift of the RGD-QWNTs after removal of surface proteins on EVs. All error bars represent standard deviation from the mean values (n = 3), except for Fig. 2F, where n = 2.

The fluorescence responses of RGD-QWNTs were investigated via incubation with the EVs in 20% human plasma. Similar to the results of integrin-specific responses, the response of the RGD-QWNTs saturated within 60 min of incubation (**Fig. 4C**). The 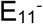 emission of RGD-QWNTs quantitatively responded to increasing concentrations of all three EV types from 10^4^ to 10^8^ EVs·mL^-1^ (**Fig. 4D**, S16). When 1×10^8^ EVs·mL^-1^ was added to the solution, the 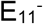 emission band of RGD-QWNTs blue shifted by 1.6 nm while the E_11_ emission remained unchanged (**Figs. 4E**, S17). The limits of detection of RGD-QWNTs for EVs were determined to be 2.4×10^4^, 2.7×10^4^, and 4.1×10^4^ EVs·mL^-1^ for HDFa, A172, and OVCAR4, respectively. No spectral shifts were observed when the same tests were conducted using QWNTs without RGD conjugation (Fig. S18). These results indicate that tripeptide-conjugated quantum well defects are the molecular focal point of interactions between nanotubes and EVs.

To assess the effects of surface proteins on the sensor responses, we compared intact and trypsin-treated (“shaved”) EVs (Fig. S19). Mild treatment of trypsin (0.01% for an hour at 37 °C) shaved surface proteins on EVs while preserving their structural and morphological integrity, as confirmed by NTA and TEM (Figs. S20-21). The supernatant from the shaved EV preparation, containing cleaved surface proteins, was further analyzed via immunoassay (**Fig. 4F**). We compared the concentrations of integrin αV in the supernatant collected after trypsin treatment of intact and protein-shaved EVs. Integrin αV was detected in the supernatant of intact HDFa-, A172-, and OVCAR4-derived EVs. In contrast, the supernatant of shaved EVs showed integrin levels below the instrument detection limit. Notably, the addition of shaved EVs no longer induced the fluorescence response of RGD-QWNTs (**Fig. 4G**, S22). The TEM images of the mixture of RGD-QWNTs and EVs indicated physical interaction without disrupting the EV structures, yet no corresponding spectral response was observed (Fig. S23). The results suggest that the RGD-QWNT response requires surface integrin on the EV and is abolished when integrin is removed, even when the EVs remain structurally intact. This establishes integrin recognition as the molecular event driving the sensor response, consistent with the specific RGD engagement and localized defect-site perturbation identified by MD simulations of **Fig 3**.

### Surfaceome protein profiling of extracellular vesicles

Our sensor unexpectedly detected fibroblast-derived EVs with response magnitudes comparable to cancer-derived EVs (**Fig. 4**), suggesting that RGD-receptor integrins are present at functionally relevant levels on healthy fibroblast EV surfaces. This possibility was not directly addressed in prior EV integrin proteomics studies, which have largely focused on cancer-derived vesicles^23^. To test this, we isolated EV transmembrane proteins and their extracellular interactors (see Methods) and performed mass spectrometry-based proteomics analysis on the EV surfaceome. We identified 1318 total proteins across the three EV types, including 125 of which were EV surface membrane proteins, of which 7 were commonly shared (**Fig. 5A**). When considering both the transmembrane surfaceome and the extravesicular interactors, the overlap expanded to 306 commonly shared proteins out of the total of 1,318 identified (Fig. S24).

**Figure 5.**
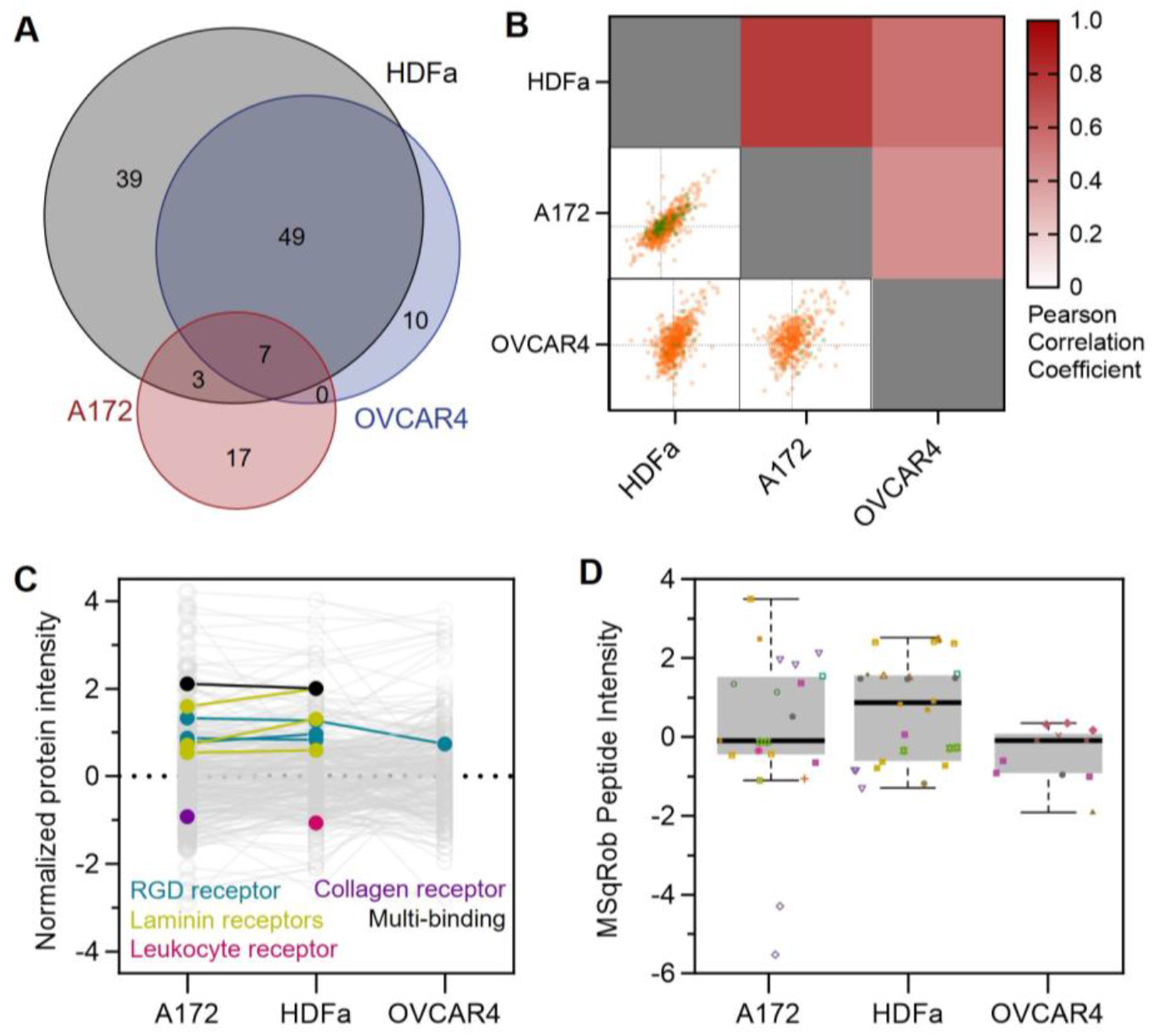
Surfaceome analysis of extracellular vesicles. (**A**) Venn diagram of detected transmembrane proteins from HDFa, A172, and OVCAR4-derived EVs. (**B**) Pairwise Pearson correlation of protein abundance. Lower triangle: log-transformed intensities of transmembrane proteins (orange) and extracellular interactors (green); Upper triangle: correlation coefficient. (**C**) Normalized abundance profiles of transmembrane proteins and extracellular interactors. Each gray point represents an individual protein; colored points highlight integrins classified by ligand-binding specificity (black, multi-binding; yellow, laminin; blue, RGD; pink, leukocyte; purple, collagen receptor). (**D**) MSqRob peptide level intensity distributions for integrin αV. Different colors denote distinct integrin αV peptides identified across EV subpopulations. Despite distinct overall surfaceome compositions among the three EV types, as confirmed by PCA clustering (Fig. S25) and low-to-moderate Pearson correlations of z-score normalized protein intensities, RGD-receptor integrins were consistently identified among the higher-abundance proteins in all three populations (**Figs. 5B**, S26–27). This shared integrin abundance, despite divergent broader surfaceomes, motivated quantitative comparison of integrin levels across cell types.

Rank-ordered intensity analysis of transmembrane proteins and extracellular interactors showed that integrin α5, αV, and β5 were abundant surface proteins across all three EV types (**Fig. 5C**, S28). These three subunits which form the RGD-binding heterodimers (α5β1, αVβ3, αVβ5), each of which engages RGD motifs via MIDAS-coordinated mechanisms.^16^ Relative protein abundance and peptide-level intensity analyses further confirmed the enrichment of these integrins in HDFa, A172, and OVCAR4 EVs, while their relative proportions varied across cell types (**Fig. 5D**, S29).

## Discussions

We developed a QWNT-based nanosensor to detect integrin subtypes and extracellular vesicles in complex biological media. Quantum defect sensitization was achieved by covalent conjugation of integrin-recognition RGD tripeptides to the quantum well defects. Molecular dynamics simulation confirmed stable binding of integrin-mimetic domains to RGD-QWNTs, supporting specific interactions at quantum defects. The specific integrin binding, coupled with dynamic changes in the biomolecular corona near the RGD-coupled quantum defects, resulted in robust hypsochromic shifts in defect emission bands, enabling quantitative detection of OVCAR4, HDFa, and A172-derived EVs.

The nanosensor detects multiple integrin subunits--α5β1, αVβ1, and αVβ3 in this study--at sub-picomolar concentrations, which is at least four orders of magnitude more sensitive than immunoassay-based quantification (nanograms per mL range). Integrin-specific interactions enabled the robust detection of EVs at concentrations as low as 10^4^ EVs·mL^-1^ in a protein-enriched plasma environment. The sensitivity is comparable to the detection limits of existing EV sensing platforms in cleaner systems, such as buffer solutions^24-27^. The enhanced sensitivity in EV detection compared to immunoassay-based quantification can be attributed to the multivalent nature of RGD-tripeptides. The surface of EVs is enriched with various membrane proteins, including multiple integrin subtypes^3^, that can serve as multiple binding sites for the nanosensor. The multivalency presumably enhances overall binding avidity, resulting in sustained, substantial sensor responses, even at low EV concentrations.

Molecular dynamics simulations provide mechanistic insight into the selective 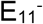 response of RGD-QWNTs upon integrin binding. The RGD-coupled quantum well defect directly docks within the integrin binding pocket and remains stably engaged throughout the trajectory. This localized, persistent contact at the defect site accounts for the selective reshaping of the dielectric environment at the quantum defect and selective 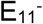 emission responses. Energetic decomposition further indicates that the interaction is dominated by electrostatic contributions with smaller but non-negligible van der Waals contributions, consistent with the charged guanidinium-carboxylate contacts that define canonical RGD-integrin recognition.^16,19^

Surfaceome proteomics connect our MD simulations to the sensor responses. The RGD receptor integrins are abundant across all three EV populations. Healthy primary dermal fibroblast EVs carry RGD-receptor integrins at levels comparable to those of cancer-derived EVs. This result indicates that integrin-targeted EV sensing generalizes beyond cancer-EV context in which EV integrins have been characterized as drivers of organotropic metastasis^23,28^ and proposed as circulating biomarkers^29,30^. Integrin abundance alone is insufficient to distinguish stromal-from tumor-derived EVs in complex biofluids, and orthogonal markers will be required to distinguish stromal-from tumor-derived EVs in complex biofluids.^31^

Finally, while we present a proof-of-concept of tripeptide-conjugated QWNTs were targeted exclusively for integrin-expressing EVs, the modular design can be extended to incorporate any short peptides targeting either other proteins or small molecules.^32^ Thus, this approach can be leveraged to identify disease-associated EV subpopulations^4,33^ and to monitor dynamic changes in EV surface composition in response to cellular stress or therapeutic perturbation in biological media.^25^

## Methods

### Synthesis of quantum well nanotubes (QWNTs)

QWNTs were synthesized using a previously reported solid-state functionalization approach.^7^ Briefly, the aqueous solution of 4-aminobenzoic acid (10 mg·mL^-1^, Sigma Aldrich) was mixed with 1.1 equiv. of nitrosonium tetrafluoroborate (Sigma Aldrich). After 2 minutes of incubation, 1 mL of the solution was added to 100 mg of raw SWCNTs (SG65i, Sigma Aldrich) in 10x PBS (pH 7.4). The mixture was vortexed for 10 s prior to the addition of 50% w/v hydrogen peroxide (Fisher Scientific), followed by 30 min incubation protected from light. The nanotubes were thoroughly washed with water and ethanol by vacuum filtration through a 20 µm polyethylene frit, then dried in an oven at 200 °C for 2 hours.

### Bioconjugation of RGD to QWNTs

0.2 mg of QWNTs was dispersed in 800 µL of 1X PBS (pH 6.0). A 100 µL portion of EDC (3.7 mg·mL^-1^, 20 mM, Sigma Aldrich) and a 100 µL portion of NHS (5.7 mg·mL^-1^, 50 mM, Sigma Aldrich) were added to the QWNTs over 30 min at 25 °C with vigorous stirring. The reaction mixture was washed with DI water three times using 5 mL syringes fitted with 20 µm polyethylene frits. NHS-activated QWNTs were dispersed in 800 uL of 1X PBS (pH 7.4). A 200 µL portion of Arg-Gly-Asp (RGD, 7 mg·mL^-1^, 20 mM, Biomatik) was added to the NHS-activated QWNTs over 2 h at 25 °C with vigorous stirring. The reaction mixture was washed with DI water three times by 5 mL syringes fitted with 20 µm polyethylene frits. RGD-coupled QWNTs were dispersed in 1 mL of 10X PBS (pH 7.4). Finally, A 6 µL portion of ethanolamine (92.9 µmol, Fisher Scientific) was added to the RGD-coupled QWNTs over 30 min at room temperature with vigorous stirring. The reaction mixture was washed with DI water three times using 5 mL syringes fitted with 20 µm polyethylene frits.

### Aqueous dispersion of carbon nanotubes

0.2 mg of carbon nanotubes (either raw SG65i, QWNTs, or RGD-QWNTs) was added to 1 mL of an aqueous solution of (GT)_15_ in water (1 mg·mL^-1^, Integrated DNA Technologies) or 1 wt/v% sodium deoxycholate (Sigma Aldrich). The mixture was sonicated using a probe-tip sonicator at 60% amplitude (FB120, Fisher Scientific) at 4 °C for an hour. The resulting solution was centrifuged at 21,300 g for 30 min. The top 80% supernatant was collected and dialyzed against 1X PBS (pH 7.4) to reduce free DNA in the solution by at least a factor of 9.

### Spectroscopic characterization

For the absorption and fluorescence measurements, the nanotubes are dispersed in an aqueous solution of ssDNA or sodium deoxycholate as described in the earlier paragraph. Absorbance spectra of nanotubes were collected by a V-780 UV-Visible/NIR spectrophotometer (Jasco, Inc.). The optical density at (6,5) E_11_ was adjusted to 0.4 to avoid inner filter effects in the fluorescence measurements. Fluorescence emission spectra of nanotubes were acquired by an InGaAs spectrometer (IMA-SWIR, Phonon. etc.). The samples were excited with a 577 nm laser power supply (500 mW, Optoelectronics Tech., Co., Ltd). The light path was shaped and fed into the back of an inverted IX73P2F microscope (Olympus), where it passed through an 850 nm long-pass emission filter, a 20x/0.45 IR objective (Olympus), and illuminated the samples in a 384-well clear flat-bottom UV-transparent microplate (Corning). Following the acquisition, the data were processed using custom MATLAB code that applied the aforementioned spectral corrections and background subtraction, and the fluorescence emission peaks were fitted with Lorentzian functions.

The excitation-emission fluorescence maps of nanotubes were obtained using a NS super (Applied NanoFluorecence, Houston, TX). The nanotubes were excited with light from a supercontinuum laser source, filtered through a tunable filter system. Excitation scans were performed from 500 to 650 nm with a 5 nm step size and a 10 nm slit width. Emission spectra were recorded with a 500 ms integration time at each excitation wavelength, and each spectrum was averaged over three times. SWIR fluorescence emission was collected at a 180° angle from excitation. The resulting emission spectra were measured by a 1D InGaAs detector cooled to -15°C.

Raman spectra of solid-state carbon nanotubes prior to aqueous dispersion were measured using an i-Raman® Prime 532H under 532 nm excitation at 100% power and a 500 ms integration time. The X-ray photoelectron spectroscopy of carbon nanotubes was performed using NEXSA G2 XPS (Thermo Fisher Scientific). High-resolution spectra were acquired for C 1s, N 1s, and O 1s regions.

### System preparation, geometry optimization, and force-field parameterization for Molecular Dynamics (MD) simulations

To evaluate the enhanced binding performance of RGD-QWNTs, two systems consisting of integrin and either RGD-QWNTs or control QWNTs were prepared. Integrin αVβ3 was selected as the model receptor due to the availability of its RGD-bound crystal structure (PDB ID: 1L5G).^16^ To reduce the computational cost, only the β-propeller domain (residues 1–438) and the βA domain (residues 110–353) were retained. To immobilize the integrin, harmonic position restraints (k = 5 kcal mol^-1^ Å^-2^) were applied to the alpha-carbon atoms within 10 Å of the terminals of these domains. Mn^2+^ ions were substituted with Mg^2+^, which are more physiologically relevant and for which well-refined force-field parameters are available. (6,5) QWNTs with a length of 80 Å were generated using the VMD Nanotube Builder and hydrogen-capped at both ends.^34^ Geometry optimization of the QWNTs was performed at the HF/6-31G(d) level using an implicit water solvent model, followed by electrostatic potential calculations in a vacuum at the HF/6-31G(d) level.^35^ Partial charges were then derived via RESP-A1 fitting using the R.E.D. server^36^.

All simulation systems were constructed using AmberTools^37^. Force-field parameters for the QWNTs were assigned based on GAFF^38^, referencing the specific parameters reported by Yahyavi et al.^39^. The integrin, water molecules, and ions were described using the AMBER ff19SB force field^40^, the TIP3P water model^41^, and Joung-Cheatham parameters^42^, respectively. A Langevin middle integrator with a friction coefficient of 1.0 ps^-1^ was used to integrate the equations of motion and control the temperature. A Monte Carlo barostat with an update frequency of 25 steps was used to control the pressure (1 atm). The SHAKE algorithm was employed to constrain bonds involving hydrogen atoms, and the SETTLE algorithm was used to treat water molecules as rigid. Long-range electrostatic interactions were handled with the Particle Mesh Ewald method with a cutoff distance of 12 Å. Short-range van der Waals interactions were treated by the Lennard-Jones potential and Lorentz-Berthelot combining rules with an energy switching function that ramps the energy smoothly between an inner cutoff of 10 Å and an outer cutoff of 12 Å. Periodic boundary conditions were imposed to all directions of the simulation box. All simulations were performed with the OpenMM package (version 8.1.1).^43^

### Equilibrium MD simulations

The linear RGD motif of the RGD-QWNT was structurally aligned with the cyclic RGD peptide present in the 1L5G crystal structure. The control QWNTs were positioned using the identical spatial coordinates for the RGD-QWNT. The assembled complex was solvated in a 110.0 × 110.0 × 100.0 Å^3^ rectangular box and neutralized to a physiological salt concentration of 0.15 M NaCl. The energy minimization process was performed using the limited-memory BFGS algorithm. The heating process was conducted up to 300 K over 2 ns under NVT conditions with a 0.5 fs timestep, with positional restraints applied to the protein backbone (9.56 kcal mol^-1^ Å^-2^) and side-chain heavy atoms (4.78 kcal mol^-1^ Å^-2^). The equilibration process was conducted at 300 K and 1 atm for 2 ns in the NPT ensemble with a 2.0 fs timestep, with the gradual removal of these restraints. Following this preprocessing procedure, a production MD simulation was performed for 800 ns in the NPT ensemble at 300 K and 1 atm with a 2.0 fs timestep to investigate spontaneous binding dynamics. All structural and energetic analyses were performed using the trajectory from the 400–800 ns window. The root-mean-square fluctuation (RMSF), center-of-mass (COM) distances between the integrin MIDAS (Mg^2+^) and the QWD, and the number of specific hydrogen bonds were calculated using CPPTraj.^44^ Interfacial contacts and contact frequency maps were computed utilizing MDTraj,^45^ where a valid contact was defined as any integrin atom located within 3.5 Å of the QWD or the nanotube sidewall. The nonbonded interaction energy between the QWD and the integrin interface was evaluated by re-running the trajectories and calculating the isolated electrostatic and van der Waals contributions using OpenMM.

### Steered MD simulations

To select the most populated binding configuration as the initial structure for these simulations, a two-step hierarchical clustering approach was employed using CPPTraj. First, macroscopic K-means clustering was performed on the entire QWNT structure relative to the integrin binding pocket across the sampled trajectory to identify the most populated binding state. Subsequently, a highly resolved secondary clustering focusing solely on the QWD of the QWNT was conducted within the identified peak interaction windows. The specific frame exhibiting the lowest RMSD to the centroid of this refined cluster was extracted. The extracted complex was solvated in a 110.0 × 110.0 × 140.0 Å^3^ rectangular box and neutralized using a salt concentration of 0.15 M NaCl. The preprocessing procedure was identical to that used in the equilibrium MD simulations, and a steered MD simulation was subsequently performed. By constraining the COM in the xy plane and the initial tilt posture of the QWNT using harmonic centroid bond forces, only the translational movement of the QWNT along the z-axis was allowed. Taking the absolute initial z-coordinate of the QWNT COM as the reaction coordinate, the QWNT was gradually pulled away from the immobilized integrin in the z-direction up to 30 Å at 1 Å intervals. At each window, a 10 ns umbrella sampling production run was performed using a pulling force constant of 5 kcal mol^-1^ Å^-2^ to collect data. Each steered MD run for two different conditions was performed to calculate the potential of mean force via the weighted histogram analysis method^46^, and the QWD-integrin binding energies were compared. The calculations were performed at 300 K with a convergence tolerance of 10^-5^ by dividing the 3.0 nm reaction coordinate into 300 bins.

### EV production and isolation

EVs were produced from cultures of A172, OVCAR4, and HDFa cell lines (ATCC) using CELLine AD1000 adherent bioreactors (DWK Life Sciences Inc., Wheaton) as previously described (Fig. S12)^47^. Briefly, roughly 25 million cells were inoculated into the cell chamber in 15 mL of complete DMEM/F-12 (10% fetal bovine serum), and the upper chamber was filled with around 800 mL of the same media. The bioreactor was adapted from complete media to DMEM/F-12 with 10% CDM-HD (FiberCell) over the course of 4 weeks. Following adaptation, once per week, the upper media chamber was refreshed with new media, and twice per week, the conditioned media in the lower cell chamber was harvested for EV isolation. The bioreactors were maintained for several months until the dialysis membranes ruptured or sufficient EVs had been collected.

For EV isolation, the conditioned media were first centrifuged at 200 g for 5 min to pellet cells and large debris, and the resulting supernatant was centrifuged at 2,000 g for 10 min to pellet smaller debris. The resulting supernatant was then centrifuged at 20,000 g for 30 min to pellet crude large EVs (Himac CP80NX ultracentrifuge, P70AT rotor), and the resulting supernatant was ultracentrifuged at 100,000 g for 60 min to pellet crude small EVs. The crude small EVs were then further purified by pooling EV-rich fractions from a 35 nm qEV original size exclusion chromatography column (Izon Science Limited), using an automated fraction collector. The EVs were then concentrated using a 100 kDa MWCO Vivaspin 500 (Sartorius) at 10,000 g.

### Nanoparticle tracking analysis

EV concentration and size distribution were determined using a ZetaView x30 Mono. Purified EV samples were diluted in 1X PBS to meet the particle count per frame of 140-200. A camera sensitivity of 83, a frame rate of 30, and a shutter speed of 100 were used to capture particle counts at the 11 positions. Post-acquisition parameters were set to a minimum brightness of 18, maximum area of 1000, minimum area of 10, and a trace length of 9 to determine if particles within each position will be counted.

### Transmission electron microscopy

EVs were negatively stained with UranyLess (Fisher Sci, NC1120874) on Carbon Type-B, 300 mesh, Copper grids. The grid was placed on 20 µl of EV solution for 10 minutes, followed by two ultrapure water washes using the same droplet technique. The grid was blotted dry using Whatman paper between each droplet. The grid was then placed, EV side down, on a 10 µl droplet of UranyLess for 2 minutes, blotted, and then left overnight to dry. The grids were imaged on a JOEL 1400 TEM operating at 120kV.

### Immunoblotting

HDFa-, A172-, and OVCAR4-derived EVs were lysed using RIPA lysis buffer with phosphatase inhibitor. Cells were also collected and lysed using the same protocol. After lysis, cell and EV-derived protein were quantified using high-sensitivity BCA (Pierce™ BCA Protein Assay Kit, Thermo Fisher Scientific). Normalized protein across cells and EVs (1 μg per lane) were loaded and separated in an SDS-PAGE (Novex 4–20% Tris-Glycine Plus WedgeWell Gel, Thermo Fisher Scientific) at 180 V for 50 min at 4 °C. The gel was then transferred to a nitrocellulose membrane (Thermo Fisher Scientific) at 10 V for 60 min at 4 °C.

Following transfer, membranes were blocked using 5% Bovine Serum Albumin (BSA) in TBS-T for 1 h at room temperature while gently rocking. The membranes were then immunoblotted with GAPDH (Abcam ab9485, 1:1000, reducing), GRP94 (Abcam ab3674, 1:1000, reducing), and CD81 (Abcam ab79559, 1:1000, non-reducing) primary antibodies for a minimum of 12 hours at 4 °C. Next, membranes were incubated with either biotinylated anti-rabbit (Jackson Immuno Research, 111,065,046, 1:10000) or anti-mouse (Jackson Immuno Research, 115,065,071, 1:10000) secondary IgG antibodies for 1 hour at room temperature, followed lastly by HRP-conjugated streptavidin (Jackson Immuno Research, 016030084, 1:10000) for 1 h at 25 °C. The bound antibody was visualized using SuperSignal™ West Pico PLUS Chemiluminescent Substrate (Thermo Fisher Scientific) and imaged using an Azure 400 (Azure Biosystems).

### Integrin and EVs detection in spiked human plasma

Plasma-spiked PBS (35 µL) was prepared by mixing 10 µL of human plasma (final 20%) with 25 µL of 1X PBS (pH 7.4) in a 384-well plate. A 10 µL portion of RGD-QWNTs (final OD 0.4) was added to a well plate. Fluorescence spectra were measured to calculate peak shift. 5 µL of PBS, integrin, or EVs were added into the well plate at a specific concentration of interest. The fluorescence spectra of RGD-QWNTs were measured every 15 min. To evaluate selectivity, Albumin from human serum (HSA), Fibronectin (Fib), and IgG were used as control proteins instead of integrin. All measurements were conducted in triplicate for each sample.

### Limit of detection calculation

Concentration-dependent 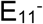 peak shifts of RGD-QWNTs in response to integrin subunits and EVs were fitted with a sigmoidal dose-response model using OriginPro 2022 (OriginLab Corporation). The LOD was calculated as formula 3.3·(SD·S^-1^), where SD and S indicate the standard deviation and the slope of the calibration curve, respectively.

### Immunoassay to quantify surface integrins in EVs

Small EVs were “shaved” with trypsin by first resuspending the crude small EV pellet after the 100,000 g spin above in 0.05% trypsin/EDTA (Gibco, 25300054) and left at 37 °C for an hour. EVs were then purified, and buffer exchange was performed by repeating the 35 nm qEV original isolation above. The EV samples were then concentrated using 100 kDa MWCO filter at 10,000 g for 15 min, with a final concentration of approximately 1×10^12^ EVs·mL^-1^. The abundance of Integrin αV on intact- and shaved-EVs was quantified using Human Integrin αV ELISA kit (Invitrogen, EH92RB) according to the manufacturer’s protocol.

### EV Proteomics

EVs were water-bath disaggregated after a freeze-thaw, as previously published, to improve accessibility of transmembrane proteins.^48^ EVs were then treated with 2 µg of PNGaseF for 2 hours at 37 °C to remove N-linked glycans, enhancing the accessibility of membrane-embedded regions for proteolytic digestion and mass spectrometry analysis. Disaggregated and deglycosylated EVs were subjected to tryptic shaving via proteomic grade proteolytic enzyme mixture (PreOmics iST DIGEST, PreOmics) at a modified volume:particle ratio of 20 µL DIGEST per 1×10^12^ particles. Tryptic peptides were desalted with the PreOmics iST kit following manufacturers protocol.

50 ng of peptides were loaded onto EvoTip Pure columns following manufacturer’s protocol, for analysis with the Evosep 1 system with a C18 PepSep Xtreme column (i.d. 75 µm, length 15 cm, 100 Å pore size, 1.5 µm particle size) connected to a timsTOF Ultra2 (Bruker, Bremen, Germany). Liquid chromatography was performed at a 450 nL·min^-1^ flow rate with a 32.5-minute gradient according to the Whisper Zoom 40 SPD method. Data was collected in positive ion mode with 100-1700 m/z mass range, 9.34 Hz ramp rate, 1.39 cycle time, 12 PASEF ramps per cycle, 100 ms ramp time, 100% duty cycle, 60 µs transfer time, and 12 µs pre pulse storage. Trapped ion mobility spectrometry (TIMS) parameters were optimized, and all subsequent data was collected using a TIMS inclusion polygon with (m/z, 1/K0) vertices of (150, 0.5), (300, 1.2), (750, 1.85), (1700, 1.85), (1700, 1.6), (1250, 1.5), (600, 1.0) and (350, 0.8) with Ion Charge Control 2.0. TIMS readouts are measured as inverse reduced ion mobility (1/K0). Ion mobility and m/z were calibrated (stdev < 0.1) with three ion peaks at m/z 622, 922, and 1221 prior to sample injections. Quality control standards were run before and after each set with iRT peptides, ensuring retention time and number of peptide identifications remained consistent throughout the run.

Data was searched using MaxQuant (v2.7.5.0) against the human FASTA (Downloaded January 2nd, 2026; 83607 entries). Oxidation of methionine and n-terminal acetylation were variable modifications, with cystein carbamidomethylation as a fixed modification. Peptide Spectral Match and Protein FDR thresholds were set to 0.01. Due to the nature of peripheral membrane domain interest, proteins identified with 1 tryptic peptide were included for analysis. However, proteins were screening for protein and top peptide probability greater than 0.9. Pre-processed data was analyzed in Perseus (v2.0.3.1) as well as via quantitative robust statistical interence software (MSqRob).^49^ Protein abundances were log2 transformed and z-score normalized using the median. Proteins were screened by 65% valid values across all samples within a group.

## Declaration of Competing Interest

M.K. is a co-founder with an equity interest in Nine Diagnostics, Inc. S.P. is an advisor with equity and intellectual property interests in Block Code Protected Ltd. The remaining authors declare no competing interests.

## Acknowledgements

This work was supported in part by the National Institutes of Health (R00-EB033580 to M.K., R00-EB033857 to C.H., R00-HL175119 to C.L.C). XPS and Raman data were acquired at the Materials Characterization Facility of the Institute for Matter and Systems at the Georgia Institute of Technology. Transmission electron microscopy imaging work made use of the BioCryo facility of Northwestern University’s Atomic and Nanoscale Characterization Experimental Center, which has received support from the Soft and Hybrid Nanotechnology Experimental Resource (NSF ECCS-2025633), the International Institute for Nanotechnology, and Northwestern’s Materials Research Science and Engineering Center (NSF DMR-2308691). I.H. was supported by the Sejong Science Fellowship Program (RS-2024-00358020) of the National Research Foundation of Korea. J.M. was supported by the Chemistry GAANN Fellowship Program. A.P. was funded through the Regenerative Engineering Program (National Institute of Health T32EB031527)

