## Supplementary Information for "Engineering Carbon Nanotube Quantum Well Defects with Recognition Tripeptides for Optical Detection of Extracellular Vesicles in Plasma"

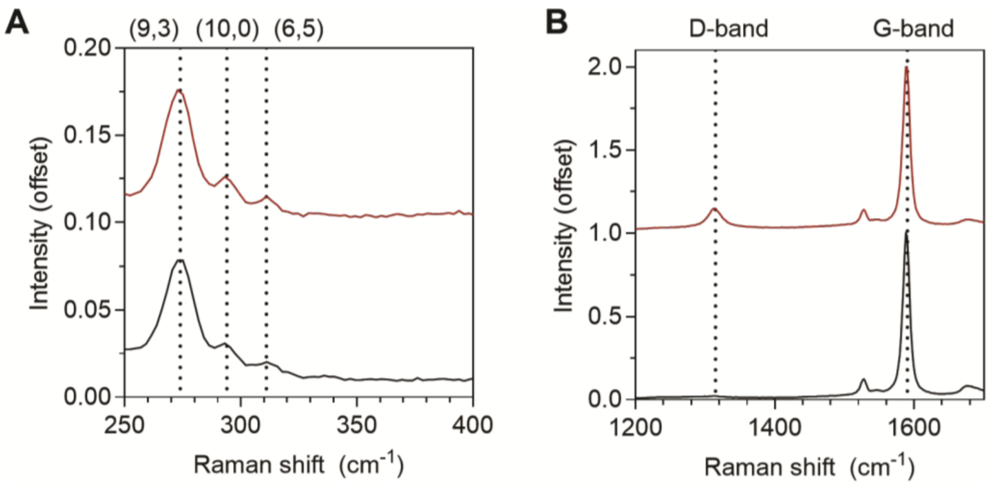

Supplementary Figure 1. Raman spectra of unfunctionalized carbon nanotubes (black) and QWNTs (red). (A) Radial breathing mode (RBM) bands of (9,3), (10,3), and (6,5) SWCNTs. (B) The D and G bands. Each spectrum was acquired using 48 mW of laser excitation at 532 nm, and normalized with respect to the G-band.

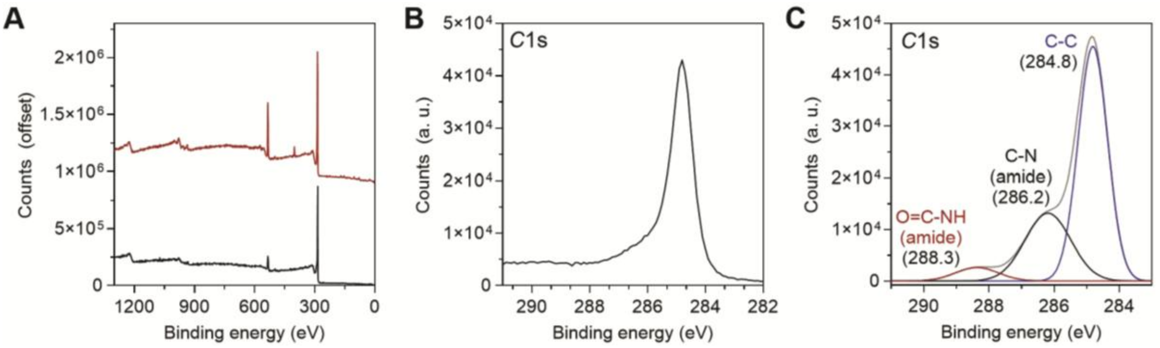

Supplementary Figure 2. XPS spectra of QWNTs (black) and RGD-QWNTs (red). (A) Full spectra of QWNTs (black) and RGD-QWNTs (red). *C* 1s XPS spectra of (B) QWNTs and (C) RGD-QWNTs.

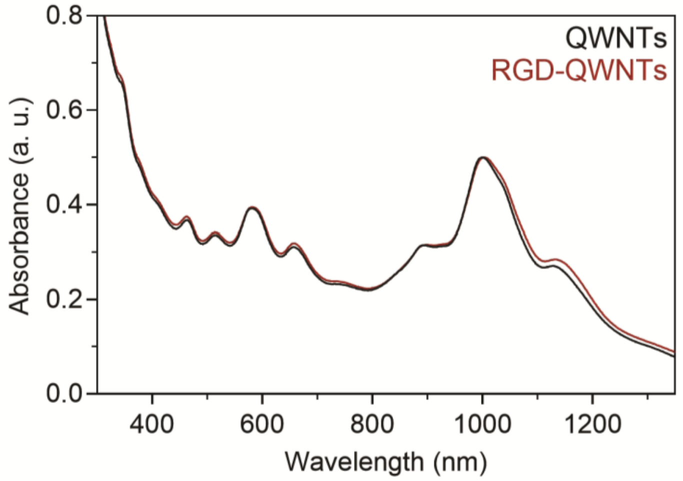

Supplementary Figure 3. Absorption spectra of $(GT)_{15}$-wrapped QWNTs (black) and RGD-QWNTs (red).

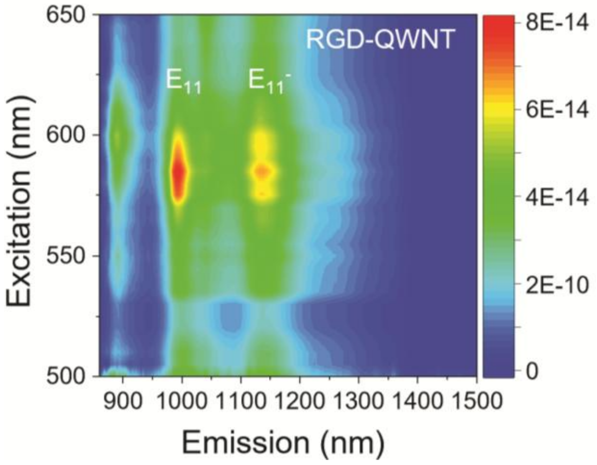

Supplementary Figure 4. Excitation-emission fluorescence map of $(GT)_{15}$ wrapped RGD-QWNTs in 1X phosphate buffered saline.

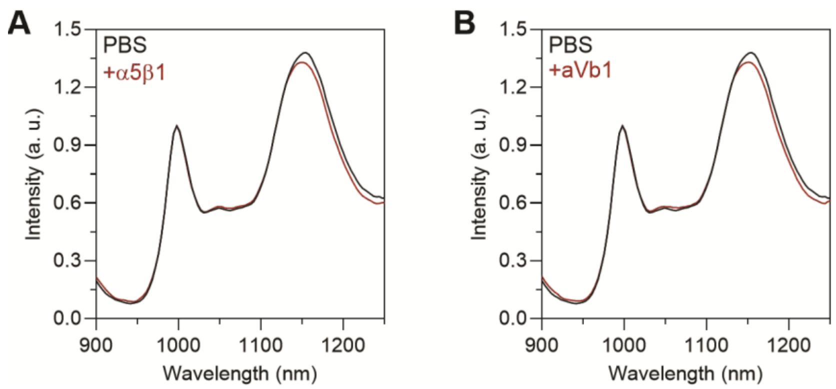

Supplementary Figure 5. Fluorescence spectra of RGD-QWNTs to integrin in 20% spiked plasma. Fluorescence spectra upon addition of PBS (black), (A) integrin α5β1, and (B) integrin αVβ1 (100 ng·mL$^{-1}$) (red).

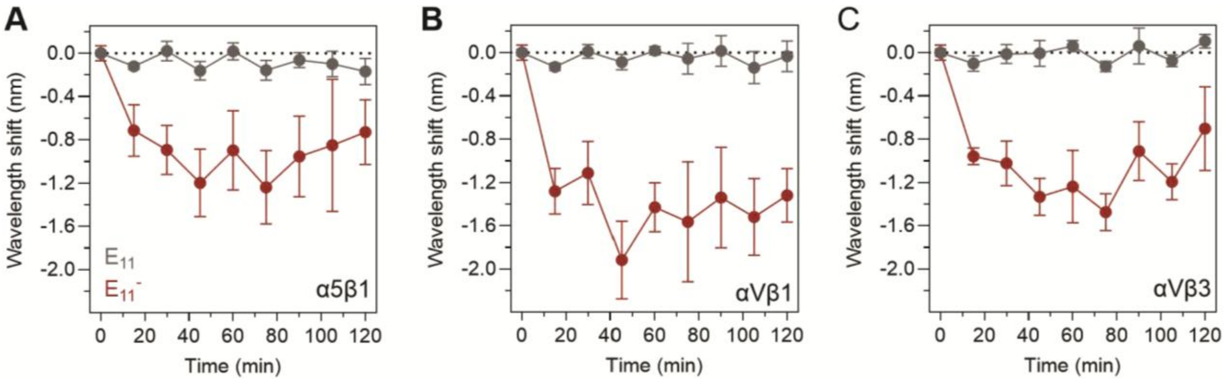

Supplementary Figure 6. Time dependent response of RGD-QWNTs to integrin in 20% spiked plasma. $E_{11}$ and $E_{11}^{-}$ peak shift of RGD-QWNTs after addition of integrin subunits (A) α5β1, (B) αVβ1, and (C) αVβ3 (100 ng·mL$^{-1}$).

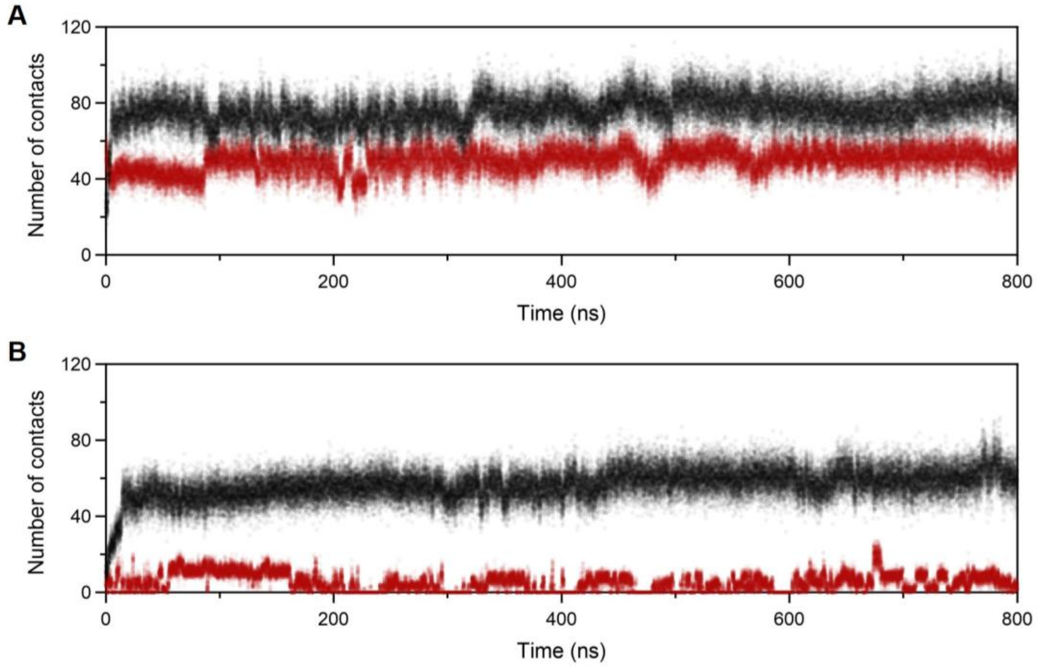

Supplementary Figure 7. Interfacial contact analysis between integrin $\alpha V\beta 3$ and QWNTs. Number of integrin atoms within 3.5 Å of the quantum well defect (red) and the nanotube sidewall (black) for (**A**) RGD-QWNT (**B**) QWNT control over the 800 ns trajectory.

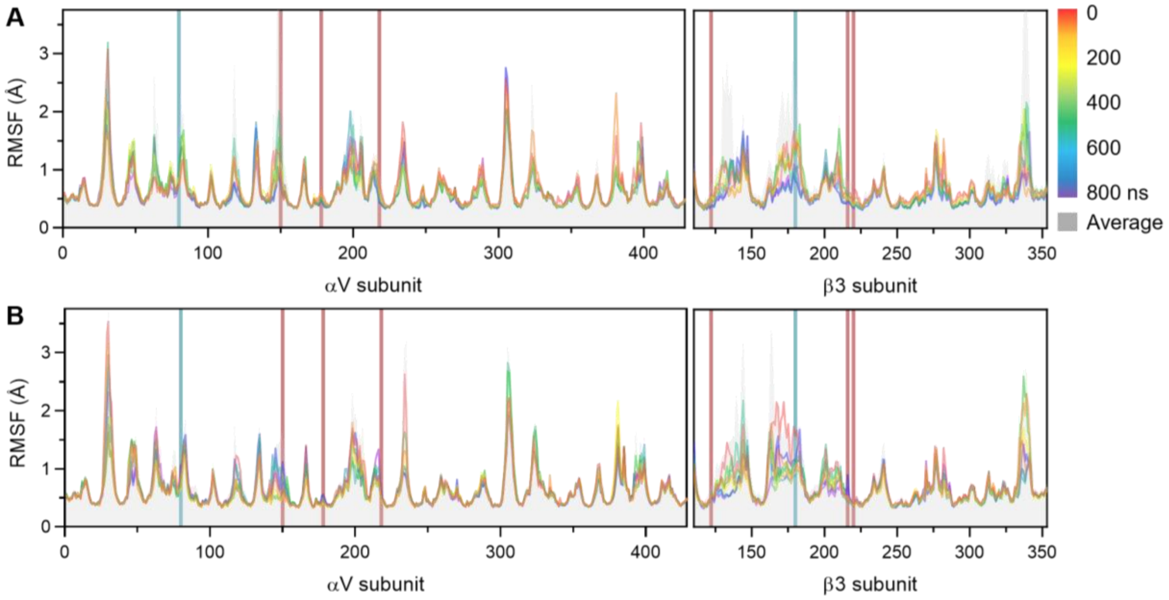

Supplementary Figure 8. Root-Mean-Square Fluctuation (RMSF) of integrin αVβ3 residues. Per-residue RMSF profiles for (**A**) RGD-QWNT and (**B**) QWNT control. Vertical lines denote known RGD-binding residues (red) and representative flexible loop positions (blue) identified in prior work by Frigerio et al (ref. 19 in the main text). Color bar represents the progression of the 800 ns trajectory.

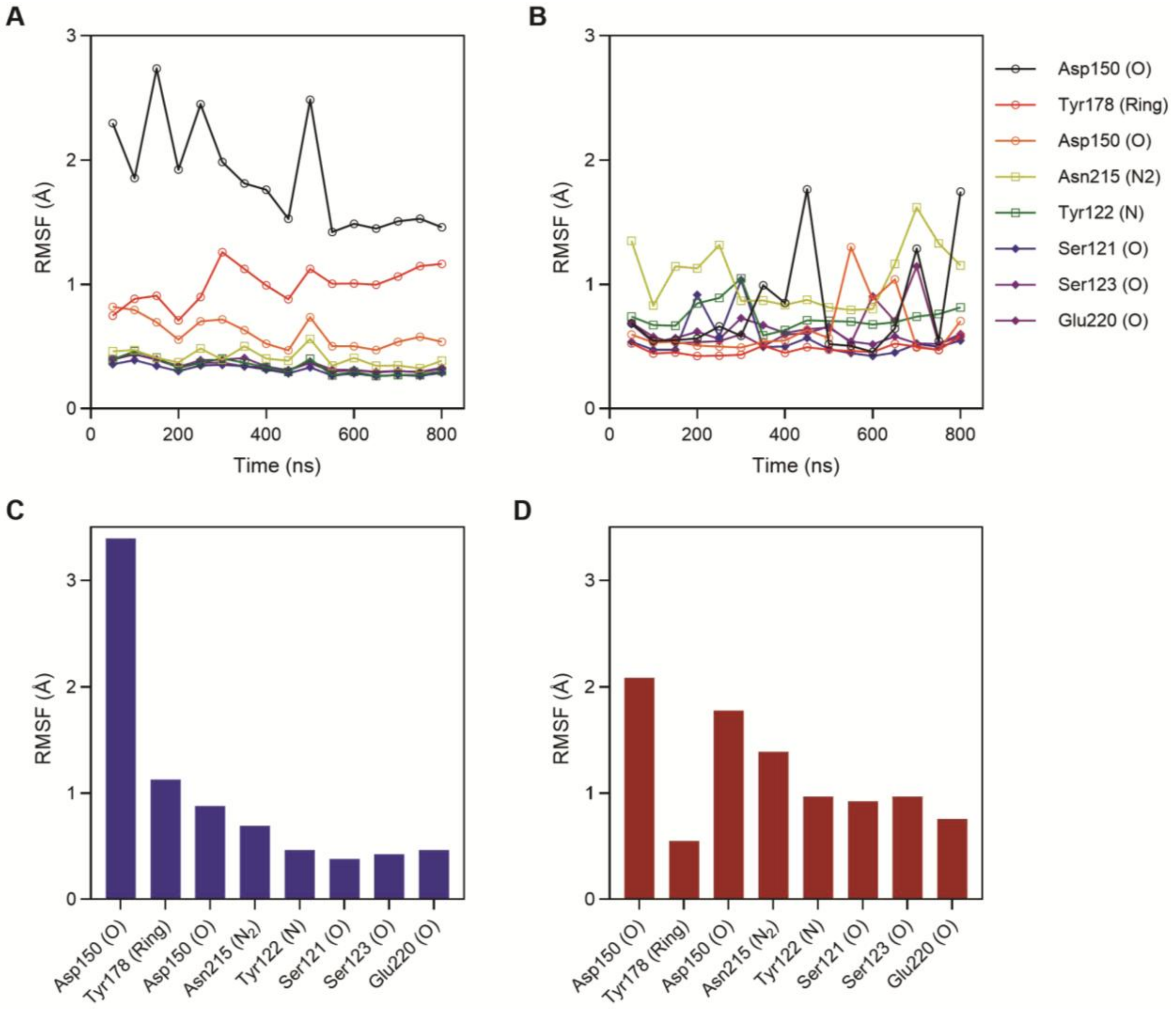

Supplementary Figure 9. RMSF analysis of RGD-binding residues at the integrin αVβ3 interface. Time-dependent RMSF profile for (**A**) RGD-QWNT and (**B**) QWNT over 800 ns. The trajectory averaged RMSF for (**C**) RGD-QWNT and (**D**) QWNT control.

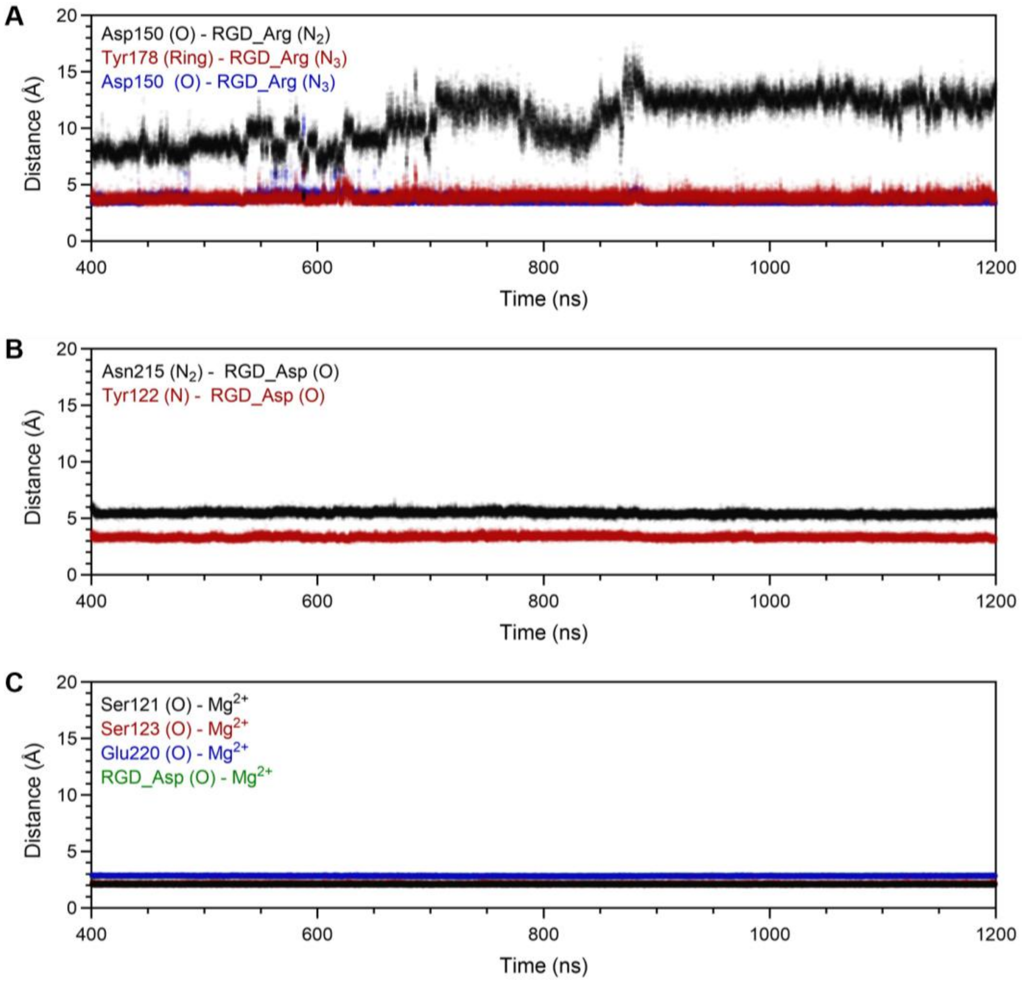

Supplementary Figure 10. Center-of-mass distance between RGD-QWNT and integrin αVβ3 residues in the (**A**) αV and (**B**) β3 subunits, and (**C**) Metal Ion-Dependent Adhesion Site ($Mg^{2+}$) in the post-equilibrium window.

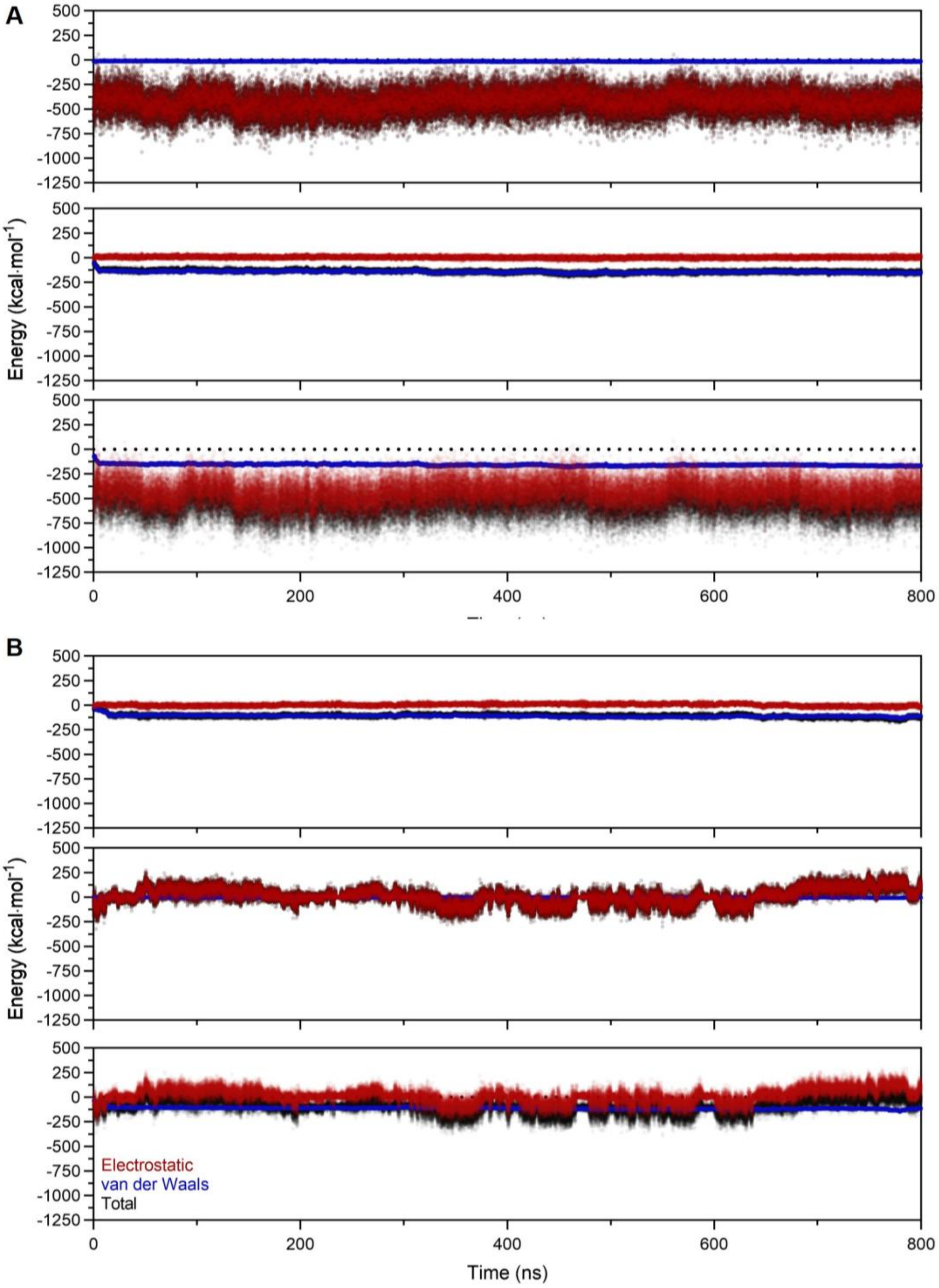

Supplementary Figure 11. Non-bonded interaction energy decomposition between integrin αVβ3 and QWNTs. Time dependent electrostatic (red), van der Waals (blue), and total (black) interaction energies for (**A**) RGD-QWNT and (**B**) QWNT control over the 800 ns production trajectory. Top: QWD-integrin interface; middle: nanotube sidewall-integrin interface; bottom: total QWNT-integrin interaction.

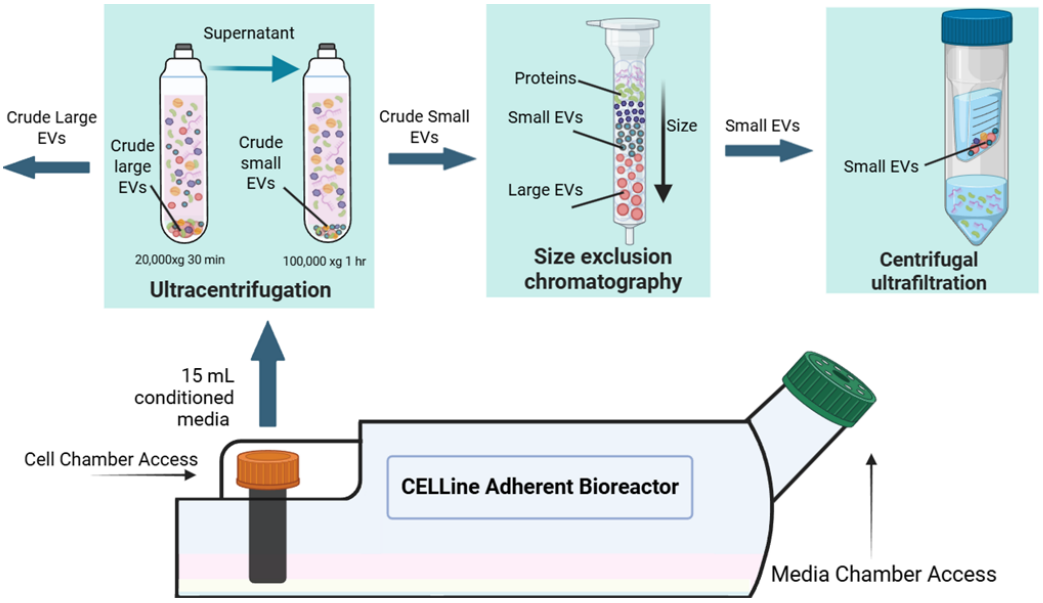

Supplementary Figure 12. Schematic illustration of EVs production and isolation.

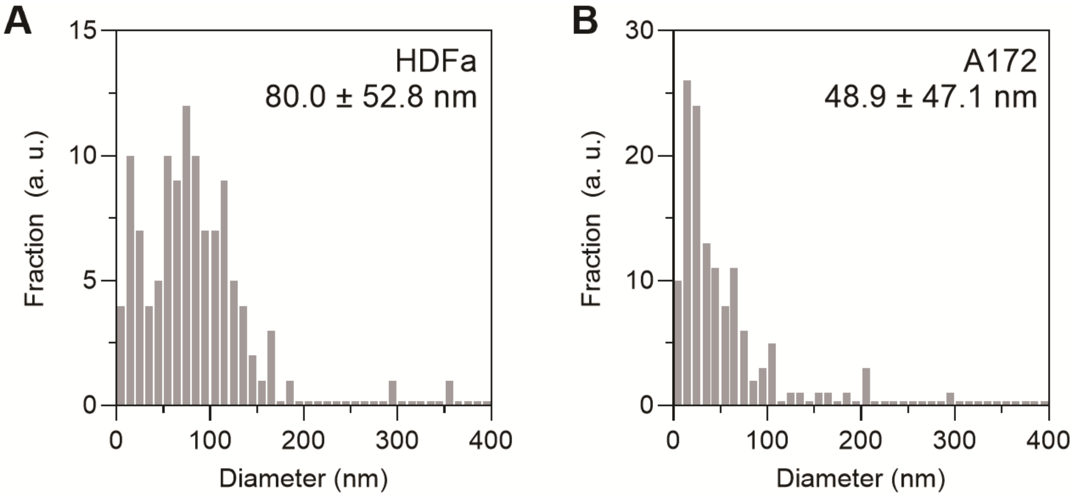

Supplementary Figure 13. Hydrodynamic size of EVs from (A) HDFa and (B) A172 cells was characterized using nanoparticle tracking analysis (NTA).

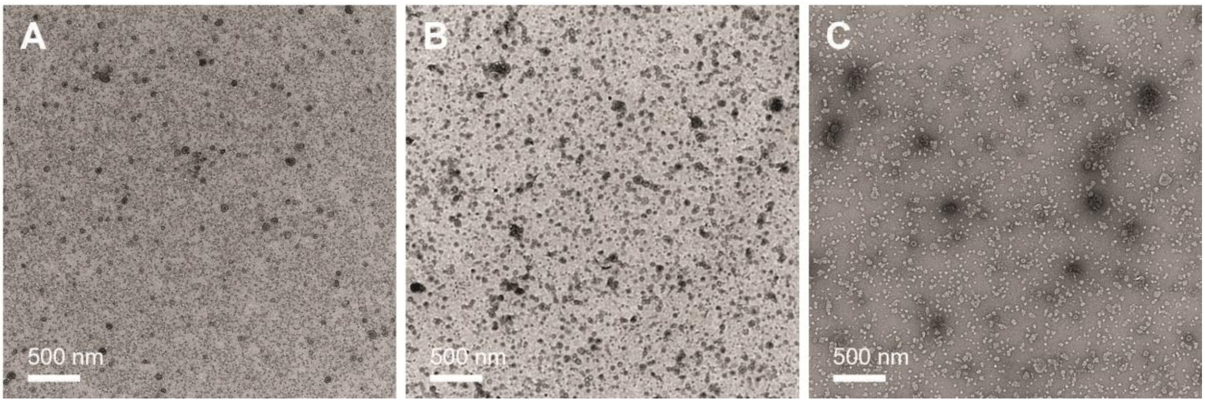

Supplementary Figure 14. TEM images of intact (A) HDFa, (B) A172, and (C) OVCAR4-derived EVs.

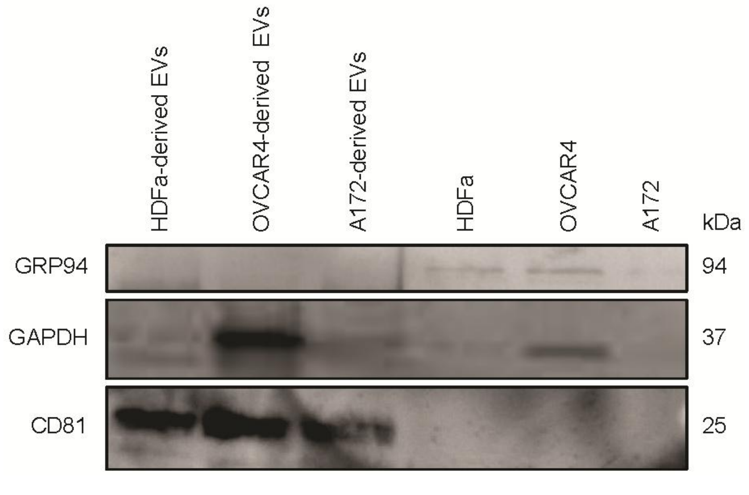

Supplementary Figure 15. Western blot of HDFa-, OVCAR4-, and A172-derived EVs and their parent cells. CD81 (EV marker), GAPDH (loading control), and GRP94 (endoplasmic reticulum marker).

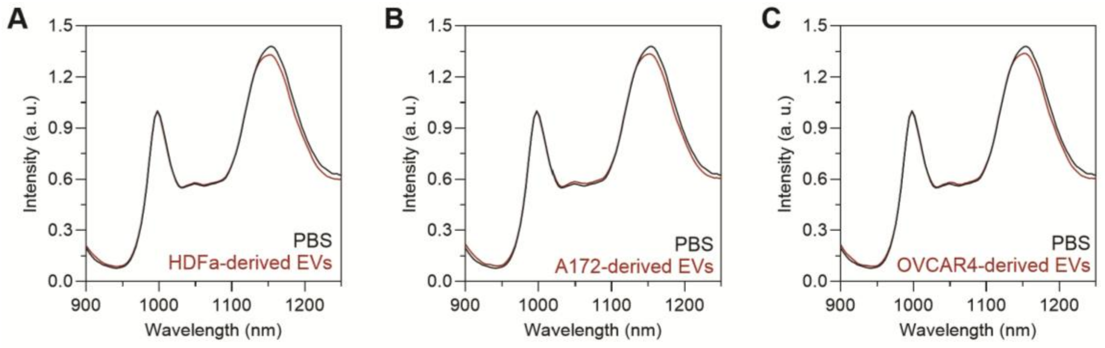

Supplementary Figure 16. RGD-QWNT fluorescence spectra to EVs in 20% plasma. Fluorescence spectra upon addition of PBS (black), (A) HDFa, (B) A172, and (C) OVCAR4-derived EVs at a concentration of 1x10$^{8}$ ·EVs mL$^{-1}$ (red). Excitation wavelength was 577 nm.

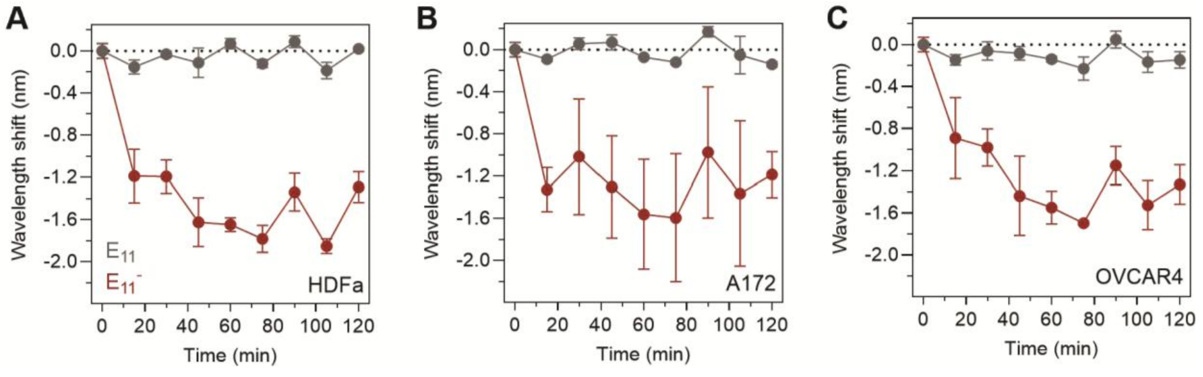

Supplementary Figure 17. Time dependent RGD-QWNT response to EVs in 20% plasma. Time-dependent $E_{11}$ and $E_{11}^{-}$ peak shift of RGD-QWNTs after addition of EVs (D) HDFa, (E) A172, and (F) OVCAR4 (1×10$^{8}$ EVs·mL$^{-1}$).

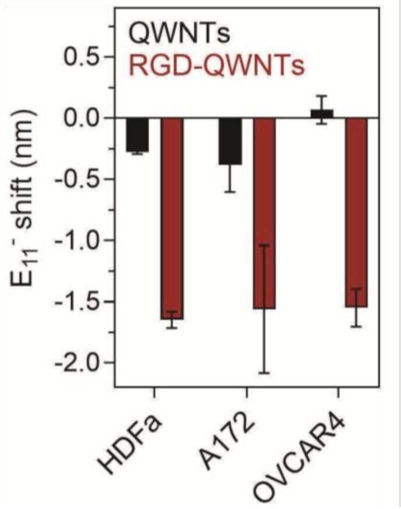

Supplementary Figure 18. QWD emission wavelength ($E_{11}^{-}$) shifts of QWNT (black) and RGD-QWNT (red) after 60 min of incubation with $1\times10^{8}$ EVs·mL$^{-1}$.

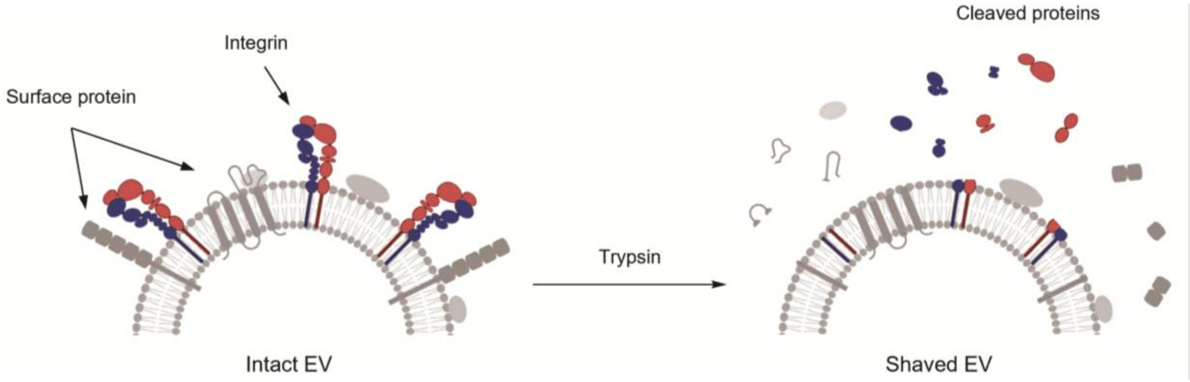

Supplementary Figure 19. Schematic illustration of surface shaving proteins on EV.

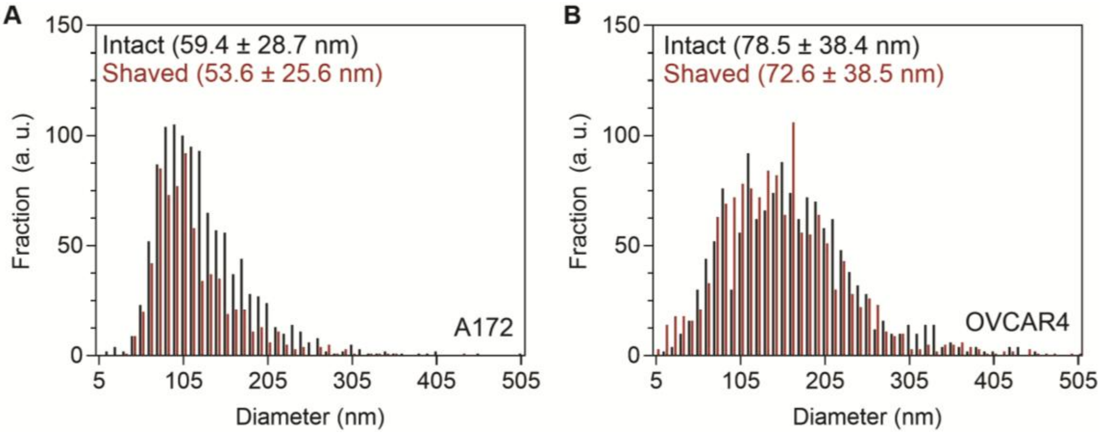

Supplementary Figure 20. Hydrodynamic size of EVs from (A) A172 and (B) OVCAR4 after trypsin treatment, characterized by nanoparticle tracking analysis.

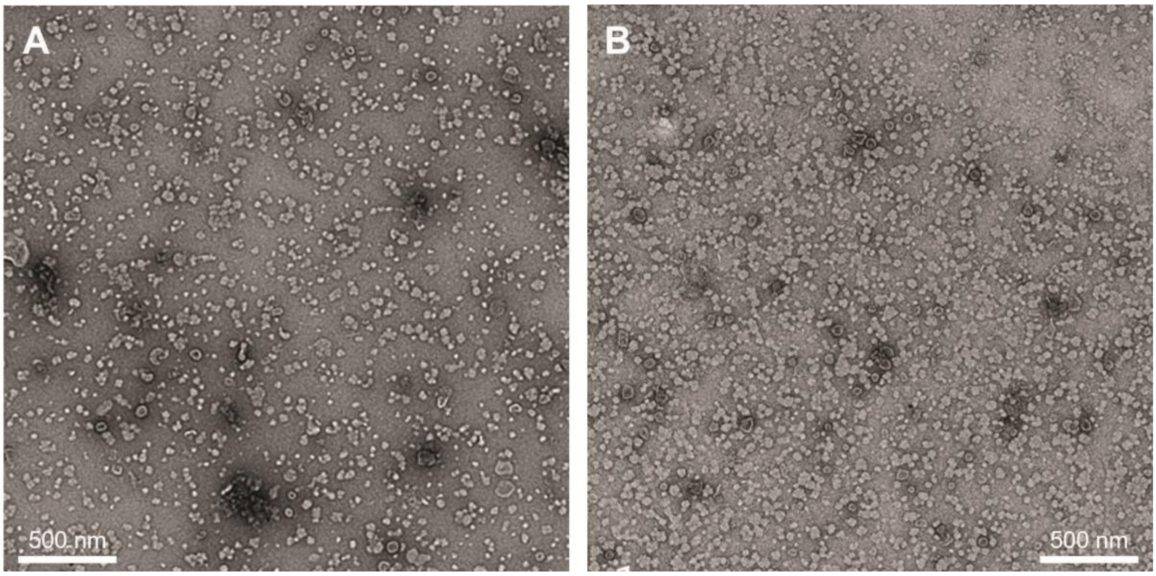

Supplementary Figure 21. TEM images of EVs after trypsin treatment. (A) intact OVCAR4, and (B) trypsin-treated OVCAR4.

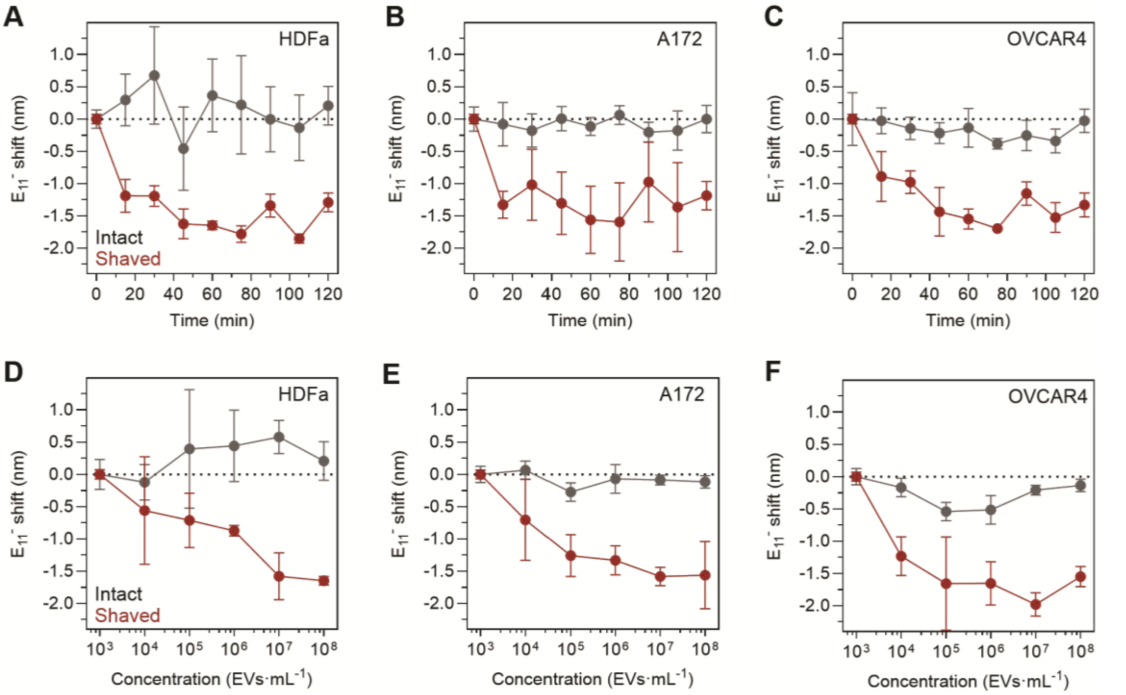

Supplementary Figure 22. RGD-QWNT fluorescence response to shaved EVs in 20% plasma. Time dependent emission peak shifts of RGD-QWNTs after addition of EVs derived from (A) HDFa, (B) A172, (C) OVCAR4 (1×108 EVs·mL-1). Concentration-dependent E11 and E11- peak shift of RGD-QWNTs after addition of EVs (D) HDFa, (C) A172, and (D) OVCAR4.

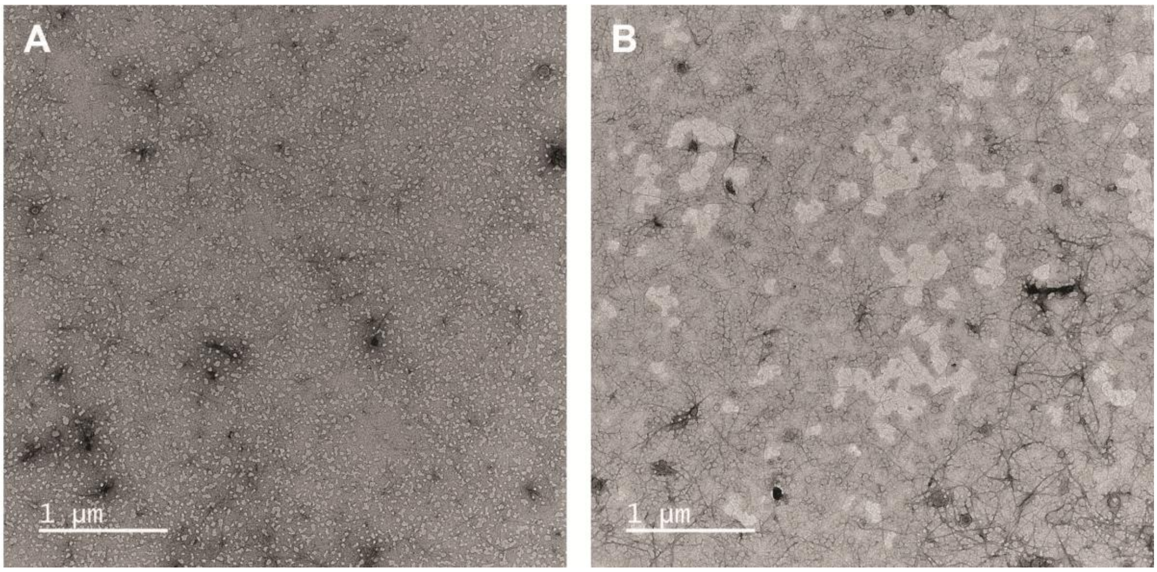

Supplementary Figure 23. TEM images of EVs after binding with RGD-QWNTs. (A) intact OVCAR4 incubated with RGD-QWNTs, and (B) trypsin-treated OVCAR4 incubated with RGD-QWNTs. White arrows indicate EVs.

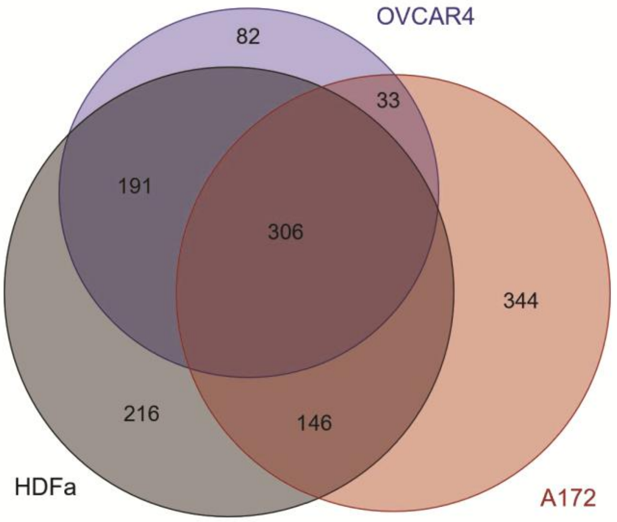

Supplementary Figure 24. Overlap of transmembrane proteins and extracellular interactors of EVs.

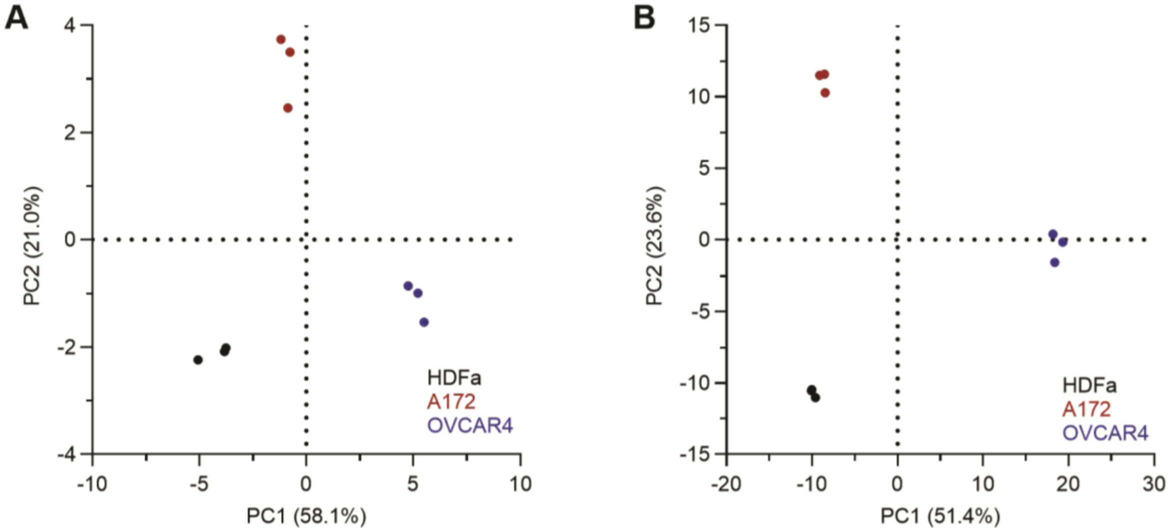

Supplementary Figure 25. Principal component analysis (PCA) of EV surfaceome proteomics. (A) transmembrane proteins and (B) transmembrane proteins and extracellular interactors across HDFa (black), A172 (red), and OVCAR4 (blue) EVs.

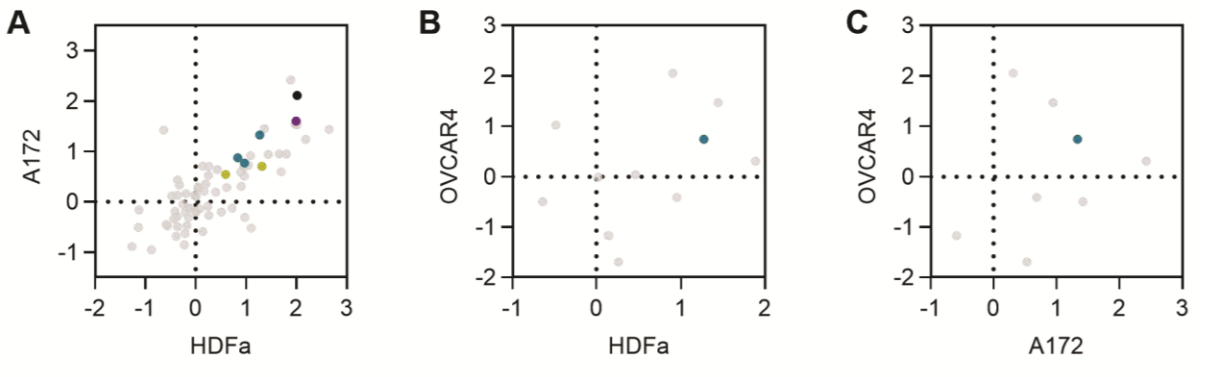

Supplementary Figure 26. Shared transmembrane proteins abundance across EV types. (A) HDFa and A172, (B) HDFa and OVCAR4, and (C) A172 and OVCAR4. Colored points denote integrin binding specificity (black, multi-binding; yellow, laminin receptor; blue, RGD receptor; purple, collagen receptor).

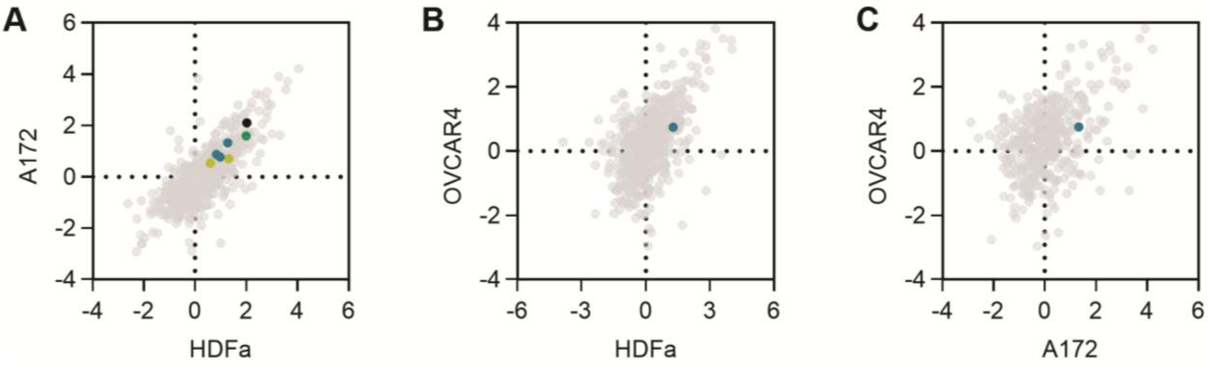

Supplementary Figure 27. Shared transmembrane proteins and extracellular interactors abundance across EV types. (A) HDFa and A172, (B) HDFa and OVCAR4, and (C) A172 and OVCAR4. Colored points denote integrin binding specificity (black, multi-binding; yellow, laminin receptor; blue, RGD receptor; purple, collagen receptor).

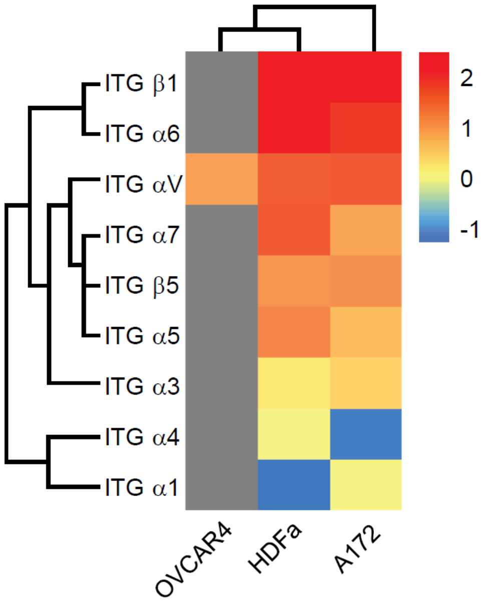

Supplementary Figure 28. Relative integrin intensity of identified in EVs. Color bar denotes the relative intensity of each integrin subtypes normalized to the total proteins identified in the EV types.

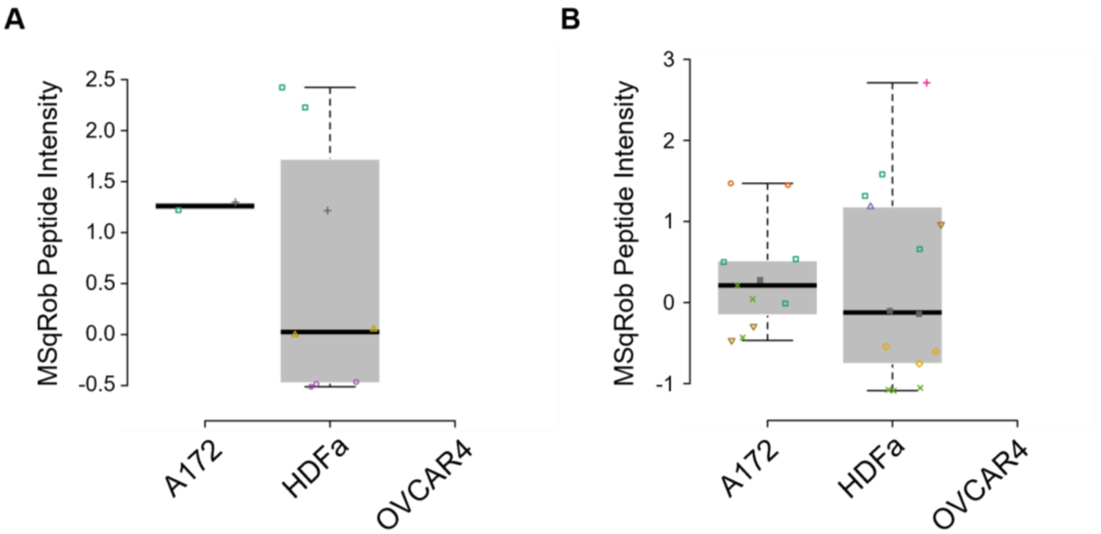

Supplementary Figure 29. MSqRob peptide-intensity boxplots for RGD-receptor (A) integrin α5 and (B) integrin β5 across A172, HDFa, and OVCAR4-derived EVs.

| System | Interaction with integrin | Electrostatic (kcal·mol$^{-1}$) | van der Waals (kcal·mol$^{-1}$) | Total (kcal·mol$^{-1}$) | Total non-bonded interaction (kcal·mol$^{-1}$) |
|---|---|---|---|---|---|
| RGD-QWNTs | QWD | −422.49 (±117.58) | −14.42 (± 4.41) | −436.91 (±117.52) | −575.23 (±117.79) |
| | Nanotube | 4.83 (± 12.38) | −143.16 (± 9.69) | −138.33 (±16.84) | |
| QWNTs | QWD | 10.54 (±78.74) | −2.75 (± 2.28) | 7.79 (±77.93) | −98.72 (± 72.20) |
| | Nanotube | 3.29 (± 14.28) | −109.80 (± 10.79) | −106.51 (±17.12) | |

Supplementary Table 1. Non-bonded interaction energies between integrin $\alpha V\beta 3$ and RGD-QWNTs and QWNTs. Electrostatic and van der Waals contributions are reported separately for quantum well defect-integrin and nanotube-integrin interactions, along with the summed total non-bonded interaction energy per system.
